# Phosphate amendment drives bloom of RNA viruses after soil wet-up

**DOI:** 10.1101/2024.10.30.616729

**Authors:** Ella T. Sieradzki, G. Michael Allen, Jeffrey A. Kimbrel, Graeme W. Nicol, Christina Hazard, Erin Nuccio, Steven J. Blazewicz, Jennifer Pett-Ridge, Gareth Trubl

## Abstract

Soil rewetting after a dry period results in a surge of activity and succession in both microbial and DNA virus communities. Less is known about the response of RNA viruses to soil rewetting—while they are highly diverse and widely distributed in soil, they remain understudied. We hypothesized that RNA viruses would show temporal succession following rewetting and that phosphate amendment would influence their trajectory, as viral proliferation may cause phosphorus limitation. Using 39 time-resolved metatranscriptomes and amplicon data, 2,190 RNA viral populations were identified across five phyla, with 37% of these predicted to infect bacteria (26%) or fungi (11%). Only 1.2% of viral populations had annotated capsid genes, suggesting most persist via intracellular replication without a free virion phase. Phosphate amendment altered RNA viral community composition within the first week and amended vs. unamended communities remained distinguishable for up to three weeks. While the overall host community remained stable, certain bacterial populations showed reduced abundance in phosphate-amended soils, likely due to increased viral lysis, as RNA bacteriophages, particularly *Leviviricetes*, proliferated significantly. Notably, 60% of the viruses with increased abundance under phosphate amendment belonged to basal *Lenarviricota* clades rather than well-known groups like *Leviviricetes*. We estimate RNA bacteriophage infections may affect 10^7^–10^9^ bacteria per gram of soil, aligning with the total bacterial population (10^7^–10^10^ g^−1^ soil), suggesting that RNA phages significantly influence bacterial communities post-wet-up, with phosphorus availability modulating this effect.

**Highlights:**

- Soil wet-up influences RNA virus dynamics for over a week
- The majority of hosts predicted for RNA viruses were bacteria and fungi
- Phosphate amendment preferentially supports *Lenarviricota* over other RNA viruses
- 60% of vOTUs that responded to phosphate were from unknown bacteriophage clades
- Most soil bacteria are predicted to be infected by RNA bacteriophages within a week

## Introduction

Soil holds the highest biological diversity of any environment on Earth (Anthony et al., 2023; Graham et al., 2024), including an extremely high diversity of viruses. Studies on soil DNA viruses consistently reveal new viruses, and soil samples taken in close proximity often contain few of the same viruses (Trubl et al., 2018, 2021; Ter Horst et al., 2021; Durham et al., 2022; Santos-Medellín et al., 2022). Soil RNA viruses are understudied, and previous studies have focused on characterising natural patterns of diversity and abundance (Starr et al., 2019; Wu et al., 2021, 2022; Chen et al., 2022; Hillary et al., 2022). These studies indicate that the diversity of RNA viruses in soil is as high or higher than that of DNA viruses, and it is shaped by environmental conditions.

While much is still unknown about viruses in soil, there is particularly little information about RNA bacteriophages (phages; RNA viruses that infect bacteria). RNA phages belong to the positive single-stranded RNA (ssRNA) class *Leviviricetes* and to the double-stranded RNA (dsRNA) family *Cystoviridae* (Walker et al., 2022; “Current ICTV taxonomy release,” n.d.); a recent survey of the global RNA virome revealed a new putative order of dsRNA phages, *Durnavirales* (Neri et al., 2022). Of the 889 species of RNA phages accepted by the international committee on taxonomy of viruses (ICTV), only eighteen genomes have been fully sequenced (https://ictv.global/taxonomy/). All of these phages infect *Proteobacteria*, with the exception of a *Cystovirus* that is thought to infect *Streptomyces avermitilis* (Callanan et al., 2018). RNA phages have generally been isolated on bacterial taxa relevant to humans, such as *Pseudomonas aureginosa* and *Escherichia coli*, and little is known about their diversity or function in natural environments.

System perturbations are considered ideal times to study the ecology of viruses, as disequilibrium is followed by a succession process involving fast growth and mortality (Teeling et al., 2012; Hahnke et al., 2015; Needham and Fuhrman, 2016). During community succession, one of the mechanisms that may drive mortality of fast-growing organisms is viral infection, following Lotka–Volterra or ‘kill-the-winner’ dynamics (Thingstad, 2000; Ignacio-Espinoza et al., 2020; Sokol et al., 2022). Wet-up is a common perturbation in soils with Mediterranean climates—the moment of first rainfall after a prolonged dry season—and leads to a disproportionate release of CO_2_ (the ‘Birch effect’; Birch, 1958). This perturbation appears to act as an induction event and causes a succession of bacteria and DNA viruses, with a substantial amount of microbial cell death attributed to viral activity (Blazewicz et al., 2020; Nicolas et al., 2023; Santos-Medellín et al., 2023).

The elemental composition of viruses is quite different from that of cells, as they are composed almost entirely of proteins and nucleic acids and are devoid of other cellular components. Thus, the stoichiometric ratio of carbon, nitrogen, and phosphorus (C:N:P) in viruses is much lower than that of cells, and it is hypothesized that a bloom of viruses may cause a local phosphorus limitation (Kuzyakov and Mason-Jones, 2018). However, viral lysis releases dissolved organic material from cells, which could potentially alleviate nutrient limitation (Tong et al., 2023). Whether or not limitations are alleviated may depend on the host and viral infection dynamics. While lytic infection (viral lysis of host cell) is one outcome of viral infection, some viruses may remain within the host cell as lysogens (via a lysogenic infection; with or without integration into the host DNA), whereas others can cause a chronic infection in which viruses are released without lysing the host (Roux et al., 2019; Lerer and Shlezinger, 2022). The main difference between these infection strategies from a macronutrient point of view is that lytic infection creates progeny viruses, which require additional nitrogen and phosphorus for protein capsids and nucleic acid, whereas lysogenic infection does not. Therefore, lytic infections may bias the C:N:P ratio more than lysogenic infections.

Here, we used time-resolved metatranscriptomics to investigate the response of RNA viruses to rewetting of seasonally dry Mediterranean grassland soil over a three-week period, with and without phosphate amendment. We hypothesized that (1) soil rewetting would cause a resurgence in the microbial community and in-turn activate the RNA virus community, and (2) phosphate amendment would increase RNA virus infection by alleviating phosphorus limitation, and thus drive changes in RNA viral community structure.

## Methods

### Soil collection and experimental setup

Soil was collected from the Buck pasture plot at the Hopland Research and Extension Center (Mendocino County, CA; 39.001767° N, 123.069733° W) at an elevation of 1,066 feet and a depth of 0–10 cm on October 3, 2020 (prior to the onset of the Autumn rains). The soil was sieved (2.0 mm wire mesh) and gravimetric soil water content measured by drying duplicate 5 g soil samples to a constant weight at 105 °C for 48 h. Soil pH was determined by adding 25 mL dH_2_O to 10 g soil followed by 1 h shaking at 400 rpm and resting for 1 h prior of measuring pH with a sensION+ pH meter (Hach) (FAO, 2021). Microcosms were established by adding 206 g (+/-0.5 g) soil to 1.9 L acid washed and autoclaved Mason jars. Microcosms were divided amongst the following treatments: (1) natural abundance water (deionized H_2_O), pH 5.5; (2) deionized H_2_O, pH 5.5 + KH_2_PO_4_; in six replicates (two treatments x six replicates x three timepoints = 36 mason jars). KH_2_PO_4_ was added to the water (for phosphate amendment treatments) and mixed for a final concentration of 50 μM. Water was added to each sample slowly and evenly across the soil surface with a syringe to bring the soil up to 30% moisture (gravimetric water content). Mason jars were sealed with lids that had airtight septa and incubated at room temperature (∼23 °C).

Microcosms were destructively sampled on day 7, 14, and 21 post wet-up. First, to measure respiration, headspace samples were collected with a 10 ml syringe that was flushed three times prior to injecting 5ml of headspace into a serum bottle. Finally, wet soil was homogenized, 5 g of soil was collected for gravimetric water content measurements and 0.5 g of soil was collected into 2.0 ml Lysing Matrix E tubes (MP Biomedicals) for DNA/RNA extractions. An additional 0.5 g of dry soil (the starting soil, not from a microcosm) was also collected into 2.0 ml Lysing Matrix E tubes (in triplicate) for DNA/RNA extraction of the starting microbial community. Extractions followed a previously published protocol ((Griffiths et al., 2000); the rest of the soil was collected and frozen at -80 °C for future analyses). Briefly, soil was combined with 200 μl TE (pH 8.0), 500 μl 5% CTAB/0.7 M NaCl/240 mM KPO_4_ (pH 8) and 500 μl of ice-cold 25:24:1 phenol/chloroform/isoamyl alcohol. Tubes were shaken in a FastPrep (MP Biomedicals Cat #116005500) for 30 s at speed 5.5 m/s, spun at 16,000 *g* at 4 °C for 5 min, and then the aqueous layer was transferred to a 2.0 ml Phase-Lock gel tube (QuantaBio Cat. #2302830). This process was repeated on the organic layer and 500 μl 24:1 chloroform/isoamyl alcohol was added to the aqueous layer and spun at 16,000 *g* at 4 °C for 5 min. The aqueous layer was added to a 2.0 ml microcentrifuge tube with 1 ml 40% w/v PEG6000/1.6 M NaCl and 1 μl GlycoBlue (ThermoFisher Cat. #AM9516) and incubated overnight at room temperature. Samples were precipitated and resuspended in 50 μl of TE (pH 8.0). The Qiagen All-Prep kit (Cat. #80284) was used to separate the RNA and DNA following the manufacturer’s protocol. The RNA samples were treated with 10 μl RQ1 RNase-Free DNase (Promega Cat. # 89836) at 37 °C for 30 min and 7 μl of 10x DNase Stop Buffer to denature the DNase at 65 °C for 10 min. DNA and RNA concentrations were quantified using Qubit 2.0 (Invitrogen), purity was assessed with a Nanodrop 2000 (ThermoFisher) (Table S1). RNA samples were shipped to Azenta for library preparation with the NEBNext Ultra II RNA Library Prep Kit (NEB, Ipswich, MA, USA), rRNA depletion with the Fast select rRNA depletion Kit (Bacteria), and sequencing. Briefly, enriched RNAs were fragmented for 15 minutes at 94 °C. First strand and second strand cDNA were subsequently synthesized. cDNA fragments were end repaired and adenylated at 3’ends, and universal adapter was ligated to cDNA fragments, followed by index addition and library enrichment with limited cycle PCR (6–14 cycles). Sequencing libraries were validated using the Agilent Tapestation 4200 (Agilent Technologies, Palo Alto, CA, USA), and quantified by using Qubit 2.0 Fluorometer (Invitrogen, Carlsbad, CA) as well as by quantitative PCR (Applied Biosystems, Carlsbad, CA, USA). The sequencing libraries were multiplexed and clustered on 1 lane of 1 flowcell. After clustering, the flowcell was loaded on the Illumina HiSeq 2500 instrument according to manufacturer’s instructions. The samples were sequenced using a 2×150 Paired End (PE) configuration, resulting in ∼350 M raw paired end reads (∼105 GB sequencing).

### Respiration measurements

CO_2_ from microcosm headspace samples was quantified with a LI-850 Gas Analyzer (Licor, Lincoln, NE), with room air as a control. A standard curve for CO_2_ was produced using the following CO_2_ concentration standards: 100 ppm, 500 ppm, 1000 ppm, and 1%. Raw data was exported from the Li-850 software and analyzed using a custom R script (version 4.4.1; https://doi.org/10.5281/zenodo.13875112).

### Metatranscriptome processing

Raw reads were trimmed with bbduk (Bushnell, 2014), removing any kmers from the negative control, mostly Illumina barcodes, primers and adapters, and also some phiX with parameters set to ftl = 5, ktrim = r, k = 23, mink = 11, hdist = 1, tpe, tbo, and minlen = 50. Bbduk was used to remove any reads containing contamination and their paired reads (parameters k = 31, hdist = 1, minlen = 50), and for read trimming (we required a minimum quality score of 10; qtrim = r, trimq = 10, minlen = 50). Following quality control, the reads were assembled using rnaviralSPAdes version 3.15.4 (Prjibelski et al., 2020). Only contigs at least 600 bp long were kept for further analysis.

### Identification of RNA viruses

Prodigal version 2.6.3 was used in meta mode to identify and translate open reading frames (ORFs) in assembled contigs (Hyatt et al., 2010). We used four sets of Hidden Markov Models (HMMs) of the RNA-dependent RNA polymerase (RdRp) protein to identify RNA viruses in the translated ORFs: (1) those from Starr et al. (Starr et al., 2019) (2) RdRp-scan (Charon et al., 2022), (3) Viral-RdRp (Olendraite et al., 2023) and (4) NeoRdRp (Sakaguchi et al., 2022). These HMMs were compared to assembled transcripts using HMMsearch in HMMer3 (version 3.3.2, http://hmmer.org/), requiring a bit score of at least 70 (Neri et al., 2022). All contigs identified by any of these HMM sets were concatenated and clustered at 95% average nucleotide identity and 85% breadth to viral operation taxonomic units (vOTUs) with mmseqs2 version 14.7 (parameters: --min-seq-id 0.95 -c 0.85 --cov-mode 0) (Steinegger and Söding, 2017). The RdRp proteins identified in contigs were also searched with palmscan (Babaian and Edgar, 2022) to verify they are not reverse transcriptases.

Quality controlled reads were mapped to the vOTUs with BBmap (Bushnell, 2014) requiring a minimum identity of 90%. The resulting BAM files were analysed with coverM version 0.6.1 with method TPM (Woodcroft BJ. 2007, https://github.com/wwood/CoverM) to generate a coverage table of vOTUs across samples requiring a minimum covered fraction of 50% (https://github.com/wwood/CoverM).

We used DRAM (DRAM-v v1.4; (Shaffer et al., 2020)) and Pharokka v1.6.1 (Bouras et al., 2023), based on multiple functional annotation platforms, to annotate open reading frames (ORFs) within vOTUs (Chen, 2004; Steinegger and Söding, 2017; Alcock et al., 2019; McNair et al., 2019; Shaffer et al., 2020; Terzian et al., 2021; Bouras et al., 2023; Larralde and Zeller, 2023). Pharokka was run on meta mode (-m) with PHANOTATE as the gene calling mode.

### Viral taxonomy assignment

Taxonomy at the phylum level was assigned based on the HMM with the highest bit score (70 minimum bit score). Higher resolution taxonomy was achieved by adding the RdRp protein sequences to existing publicly available alignments using the online tool MAFFT --addfragments (Katoh and Frith, 2012; Neri et al., 2022). Phylum-level trees were built on the mafft online platform (Kuraku et al., 2013) using average linkage (UPGMA) based on the alignments with advanced settings: keep alignment length, strategy: auto, scoring matrix BLOSUM62. The trees were refined using the interactive tree of life (IToL) v5 (Letunic and Bork, 2021). The reference sequences that remained in the pruned tree along with RdRp protein sequences from this study were aligned with mafft version 7.5 (Katoh and Frith 2012), the alignment was trimmed with trimal version 1.2 (https://github.com/inab/trimal) using parameters -gt 0.5 -conc 0.6 and a phylogenetic tree was constructed with IQtree2. We overlaid onto this tree known hosts of reference sequences (Katoh and Frith, 2012; Neri et al., 2022) to predict hosts for vOTUs from this study.

### Viral host prediction

To predict putative hosts for vOTUs, we used iPHoP (Integrated Phage HOst Prediction) v1.3.3 with database “iPHoP_db_Aug23_rw” requiring a confidence score > 90. For vOTUs representing eukaryotic viruses, we used their proximity on the phylogenetic trees to viruses with known hosts. If a vOTU clustered in a small (≤ 10 leaves) monophyletic group with a virus with a known host, it was assigned the same general group of hosts to the vOTU (e.g., mammals, fungi). In very few cases, more than one virus with a known host clustered with a vOTU with the same requirements. When the known hosts came from different groups, they were always eukaryotes, and in these cases the host was assigned as “eukaryote”.

To model the potential number of virocells, we followed the approach previously described in Nicolas et al. (Nicolas et al., 2023) with minor amendments. First, the amount of phage RNA was calculated by multiplying the proportion of reads mapped to phage genomes by the amount of RNA (in ng) per microcosm. The average mass (in Da) of a single RNA phage genome was calculated based on curated *Leviviricetes* genomes from NCBI virus (Table S2), by using the EnCor bio RNA molecular weight calculator (https://www.encorbio.com/protocols/Nuc-MW.htm), then averaging all genome weights. The average number of phage genomes per microcosm was calculated by dividing the total mass of phage RNA by the average mass of an RNA phage. To estimate how many cells were infected, the number of phage genomes in an infected host cell (virocell) was estimated. In DNA viruses, the number of phages can be divided by the average burst size to estimate the number of infected and lysed cells. However, RNA viruses do not necessarily lyse their host, but can be transferred through direct contact between hosts, thus the concept of burst size cannot be used. Instead, a range of copy number of RNA phage genomes per virocell was estimated, ranging from hundreds to hundreds of thousands. The estimate numbers are based on: (1) the average RNA phage genome size is ca. 4000 bases (single stranded), while DNA phages have genomes that are typically 10–500 times larger, and can even be as large as 735kb (Al-Shayeb et al., 2020), and (2) as very few RNA phages have been cultured (Callanan et al., 2018), it is possible that some RNA phages do not have an extracellular phase. Thus, their life cycle may not include synthesis of capsid proteins, reducing the cellular resources required to create a new phage, similar to fungal viruses (Lerer and Shlezinger, 2022). Assuming that the average bacterium infected by an RNA phage is no different than a bacterium infected by a DNA phage, the same resources can be allocated to generate many more new RNA as opposed to DNA phages.

### Microbial community analyses

The genes coding for 16S rRNA and the nuclear ribosomal internal transcribed spacer 2 (ITS2 between 5.8S and 28S) were amplified from DNA extracted from the soil microcosms. As DNA was used first for metagenomics, only 30 out of 39 samples were available for amplicon sequencing (Table S3). However, these samples represented all time points with and without phosphate amendment. Hypervariable region v4 of the 16S rRNA gene was amplified using primers 515F-Y (515F-GTGYCAGCMGCCGCGGTAA) (Parada et al., 2016) and 826R (806R-GGACTACNVGGGTWTCTAAT) (Apprill et al., 2015), and the ITS2 was amplified using primers 5.8S_Fun and ITS4_Fun (Taylor et al., 2016).

For ITS2 amplification, each reaction included 1X Phusion High-Fidelity master mix (New England Biolabs), 200 nM primers, 3% DMSO, and 0.1 uL of 1:20 diluted DNA. The amplification process involved initial denaturation at 98 °C for 3 minutes, followed by 30 cycles of denaturation at 98 °C for 10 seconds, annealing at 55 °C for 30 seconds, extension at 72 °C for 30 seconds, and a final extension step at 72 °C for 10 minutes. For 16S rRNA gene amplification, 0.1 uL-1 of 1:20 diluted DNA was mixed with 1X Kapa HIFI Hotstart Readymix (Roche) and 200 nM primers. The amplification protocol included an initial denaturation step at 95 °C for 3 minutes, followed by 25 cycles of denaturation at 95 °C for 30 seconds, annealing at 55 °C for 30 seconds, extension at 72 °C for 30 seconds, and a final extension step at 72 °C for 5 minutes. After PCR, cleanup was performed using 0.8X (for 16S rRNA) and 0.9X (for ITS) AMPure XP Beads (Beckman Coulter). Subsequently, a secondary amplification was carried out using Kapa HIFI mastermix with Illumina XT indexes to attach barcodes to the PCR products. A final cleanup step was performed using 1.2X AMPure XP Beads, followed by pooling of samples in equimolar ratios and sequencing on the MiSeq platform using a 2×250 cycle kit with approximately 25% phiX spike in.

For 16S rRNA gene sequence analysis, quality trimming of reads was performed with Trimmomatic version 0.39 with parameters LEADING:3 TRAILING:3 SLIDINGWINDOW:4:15 MINLEN:200 (Bolger et al., 2014). A negative control was also sequenced and analyzed. The Qiime2 pipeline with dada2 denoising was used (Callahan et al., 2016; Bolyen et al., 2019). Samples were rarified to 3,411 reads, leading to the discard of two samples, one of which was the negative control for which there were no merged reads, and the other was one of three replicates of time point 1 without phosphate amendment.

For ITS2 sequence analysis, as the ITS2 region varies in length, merging reads may bias against some fungal species (Yang et al., 2018). As merging led to loss of ca. 40% of reads, only the forward reads were used to avoid loss of data. Due to the range of ITS2 sizes, the minimum length after quality trimming required was at least 100 bases rather than 200 bases, and read length was truncated to 230 bases. Illumina adapters were cut from both sides of the forward reads to avoid read through artifacts, followed by read denoising with dada2 (Callahan et al., 2016). Classification of reads with the UNITE Qiime2 version 9 including all eukaryotes (215,454 taxa) (Abarenkov et al., 2023) was not successful. Therefore, ITS2 representative sequences were searched against the NCBI ITS2 database, and accession numbers converted to taxonomy with the R taxonomizr package (Sherrill-Mix, 2023). Samples where rarified to 1,887 reads per sample.

### Statistical analysis

All analyses of the virus data were performed in R. The data was manipulated using the tidyverse package (v2.0.0) (Wickham et al., 2019), and plots were generated using ggplot2 (v3.5.1) (Wickham, 2016) and ggpubr (Kassambara, 2023). Statistical analyses, including PERMANOVA, were performed with the vegan (v2.6-8), DESeq2 (v1.44.0) and ape (v5.8) packages (Love et al., 2014; Paradis and Schliep, 2019; Oksanen et al., 2020). Additional R packages used include RColorBrewer (v1.1-3) for color-blind friendly color palettes (Neuwirth, 2022), stringi (v1.8.4) for string manipulation (Gagolewski, 2022), microshades (v1.13) (Dahl et al., 2022) and dichromat (v2.0-0.1) for extended color palettes (https://CRAN.R-project.org/package=dichromat), dunn.test (v1.3.6) for Dunn statistical testing (https://CRAN.R-project.org/package=dunn.test), pheatmap (v1.0.12) for heat maps (https://CRAN.R-project.org/package=pheatmap), and ggpmisc (v0.6.0) for ggplot extensions (https://CRAN.R-project.org/package=ggpmisc). Statistical tests, ordinations and taxonomic profiles of bacteria and eukaryotes were generated via Qiime2 (Estaki et al., 2020). A rooted 16S-rRNA tree for weighted UniFrac calculation was acquired from the Qiime2 “Moving pictures” tutorial (https://docs.qiime2.org/2024.5/tutorials/moving-pictures/). Three tree is rooted at the midpoint of the longest tip-to-tip distance in the unrooted SILVA tree.

### Data availability

Raw reads are publicly available on the NCBI short read archive (SRA) under project number PRJNA1161162. The dereplicated vOTUs set, RdRp protein sequences, R code and files needed to run the code are available at https://github.com/ellasiera/RNA_virus_wetup_2023. Microbial community amplicon sequences are available in the NCBI SRA database under project ID PRJNA1161162.

## Results

We characterized the response of the RNA viral community to rewetting of a Mediterranean grassland soil with or without phosphate amendment at multiple timepoints over three weeks. A total of 3,140 unique vOTUs with an average length of 1,303 bases (minimum 600 bases, maximum 9,631 bases) were yielded. To determine patterns in vOTU diversity, reads were mapped with ≥ 50% breadth of coverage, which reduced the number of vOTUs to 2,190. The normalised abundance (in cumulative transcripts per million, TPM), taxonomy and host prediction of the vOTUs are described in Table S4.

Soil phosphate amendment significantly reduced the Shannon diversity and Pielou’s evenness index of viral communities but did not impact the overall richness of vOTUs (species number) (Kruskal-Wallis test, *p*_*Shannon*_ < 0.001, *p*_*Pielou*_ < 0.001; Fig. 1A). Both phosphate amendment and rewetting had a significant effect on the trajectory of viral beta diversity (Fig. 1B; PERMANOVA community∼Time*Phosphate *p* = 0.001 *R*^*2*^ = 15%). A clear change in RNA virus community structure was observed between T0 (before wet-up) and all subsequent time points, and between soils amended (or not amended) with phosphate (Fig. 1B). Only small temporal effects on community structure at timepoints beyond the first week following rewetting were observed.

**Figure 1.**
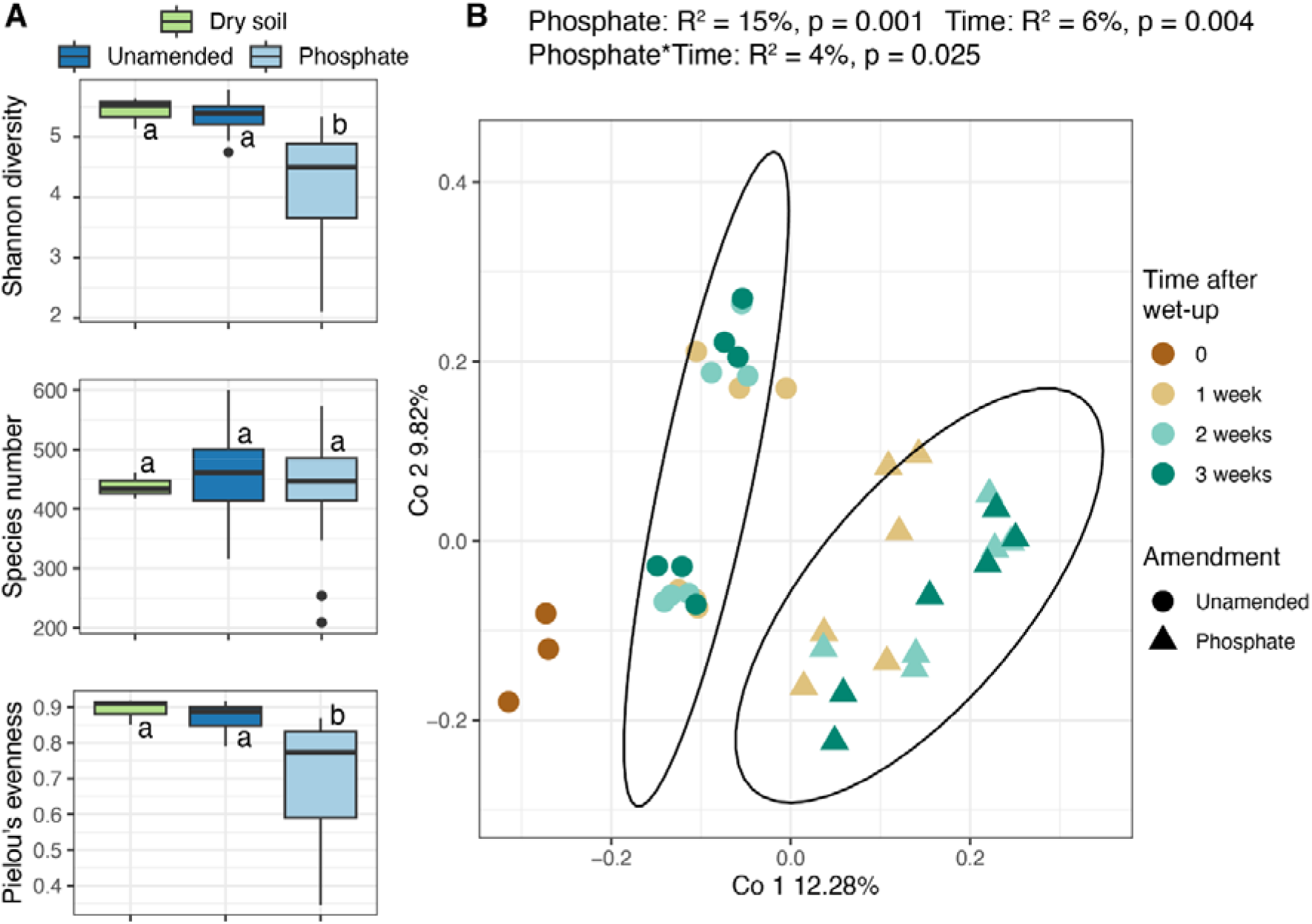
RNA virus community diversity and structure. RNA viral communities in a Mediterranean climate grassland soil before and after wet-up, and with or without phosphate amendment, characterized by (A) alpha diversity indices: Shannon’s H, richness (species numbers) and Pielou’s evenness, and (B) PCoA ordination of Bray-Curtis dissimilarity of community composition. Shapes represent presence (triangle, P+) or absence (circle, P-) of phosphate amendment. Colours represent time in weeks after wet-up. Ovals surrounding datapoints represent 95% confidence intervals. Significant treatment differences for diversity indices were determined via Kruskal-Wallis and Dunn tests with Benjamini-Hochberg correction for multiple comparisons. Community structure statistics were calculated by PERMANOVA.

We evaluated the variability in RNA virus community composition across our experiment by examining shared vOTUs in different microcosms by time and treatment. A total of 755 vOTUs appeared in only one microcosm, while 1,109 vOTUs appeared across 2– 5 microcosms, reflecting consistent rank richness across different conditions (Fig. S1, S2). At least half of the vOTUs were present in only one of three replicates, showing a substantial degree of variation between replicates (Fig. S3). Notably, 42 vOTUs appeared only in dry soil, 591 vOTUs were unique to phosphate-amended soils, and 298 vOTUs were found only in unamended soils post-wet-up (Fig. S4). 123 vOTUs were present in at least 90% of microcosms, of which the most abundant groups were *Cryppavirales* (31), basal *Lenarviricota* (17) and *Wolframvirales* (12).

To investigate putative hosts of the RNA viruses we identified, we overlaid known hosts from Neri et al. (Neri et al., 2022) onto the phylogenetic trees constructed for taxonomic assignment. Of the 2,190 vOTUs, we identified putative hosts for 807 vOTUs (37%) or roughly one third of the vOTUs identified in the study—these were further assigned as phages (68% in amended soil, 79% in unamended soil) or fungal viruses (24% in amended soil, 15% in unamended soil), with a few additional viruses infecting other eukaryotes. Of the total vOTUs with a predicted host, 560 (70%) likely infect bacteria, 184 (23%) infect fungi, 22 infect invertebrates, 15 infect plants, 17 infect unassigned eukaryotes, 4 infect mammals and 5 infect birds, reptiles, turtles or amphibians (grouped together into Sauria). Amongst the 1,533 phages identified, relatively few (27) had hosts predicted using iPHoP (Roux et al., 2023); these hosts included *Gammaproteobacteria, Bacteroidia, Bacilli, Alphaproteobacteria, Actinomycetia, Planctomycetia, Methanobacteria* and *Thermoplasmata*.

We characterized viral communities before and after wet-up by assigning RNA vOTU taxonomy based on the RdRp gene—a marker found in most RNA viruses (Edgar et al., 2022)—and then visualized the data via phylogenetic trees (Fig. S5–S9). At the phylum level, RNA viral community abundance was dominated by *Lenarviricota*, regardless of time or amendment (Fig. 2A). The *Howeltoiviricetes*, many of which infect fungi (i.e., mycoviruses), bacteriophages of the class *Leviviricetes*, and unclassified *Lenarviricota* were also abundant groups, and had the highest richness of unique vOTUs (Fig. 2B, 2C, 2D). Overall, relative abundance and richness (number of unique vOTUs) by taxonomic groups (phylum to order level) were positively correlated (*R*^*2*^ = 60%, *p* < 0.001) (Fig. 2C).

**Figure 2.**
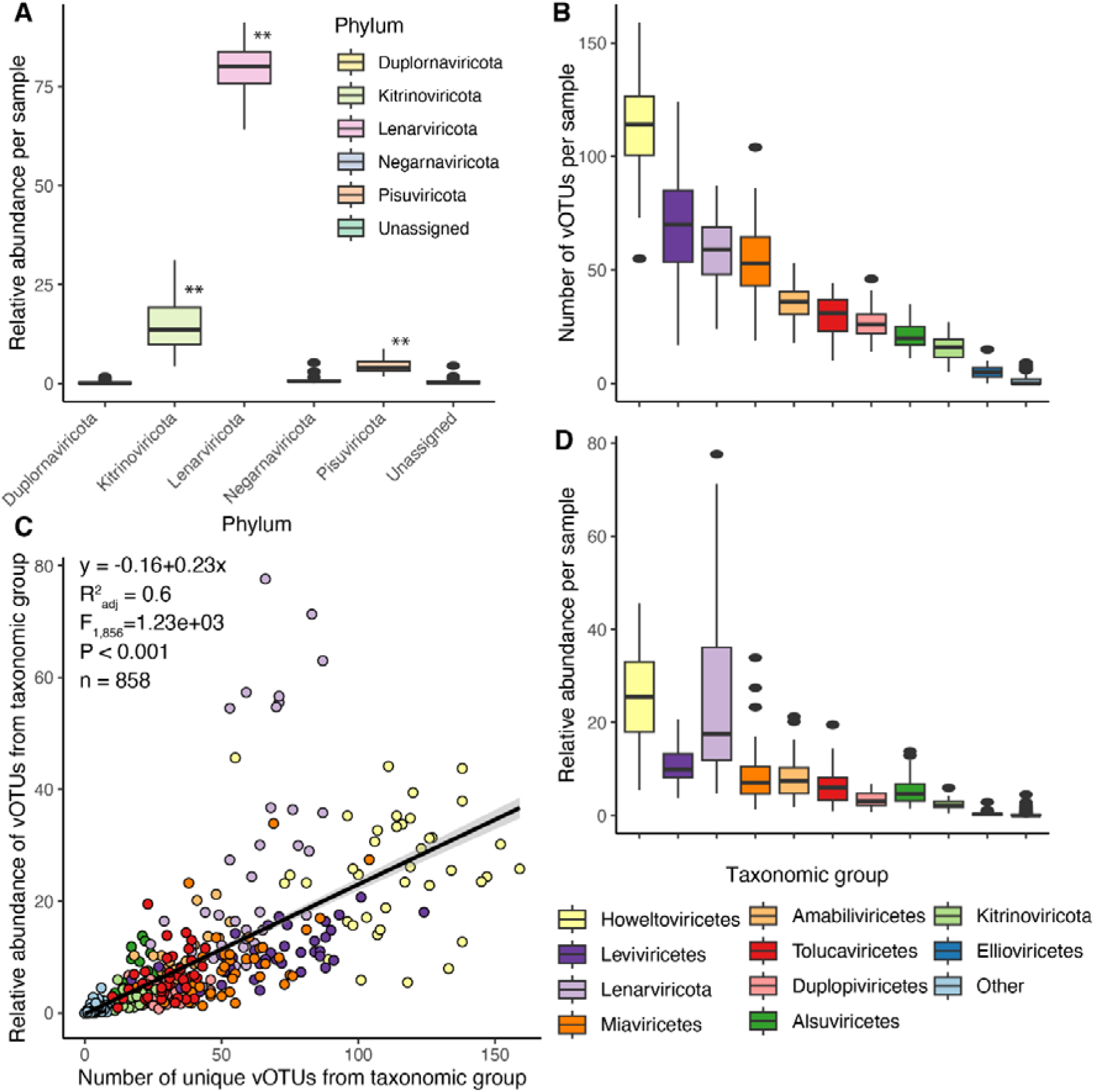
Taxonomy of RNA viral communities in dry grassland soil (T0) and after wet-up and phosphate amendment. (A) Relative abundance of different RNA virus phyla (** denotes *p* < 0.001); (B) Richness (number of unique vOTUs) per class of RNA viruses; (C) Correlation between mean relative abundance and richness of vOTUs from a specific class in all microcosms (*R*^*2*^ = 0.87, *p* < 0.001, *F* = 5140, *n* = 780). D) Relative abundance of different RNA viruses by class. The legend in panel D applies to panels B–D.

We observed a large shift in overall RNA viral community composition during the first week post wet-up, but relatively muted changes thereafter. We also investigated the temporal trajectories of specific phyla. The phylum *Lenarviricota*, which includes most phages, and the family *Narnaviridae*, which includes mostly mycoviruses, increased roughly 2-fold in abundance (regardless of phosphate amendment) during the first week after wet-up (Fig. 3A-B; Table S4). After the first week, the relative abundance of *Lenarviricota* remained constant. The second most abundant phylum, *Kitrinoviricota*, which contains many eukaryotic viruses (Wolf et al., 2018), also increased in the first week after the wet-up when phosphate was not added (Fig. S10). A similar pattern was observed in the phylum *Pisuviricota* which includes order *Durnavirales* that is thought to include phages in addition to eukaryotic viruses (Neri et al., 2022) (Fig. S10). Phylum *Negarnaviricota* (eukaryotic viruses) decreased over time regardless of phosphate amendment (Fig. S10).

**Figure 3.**
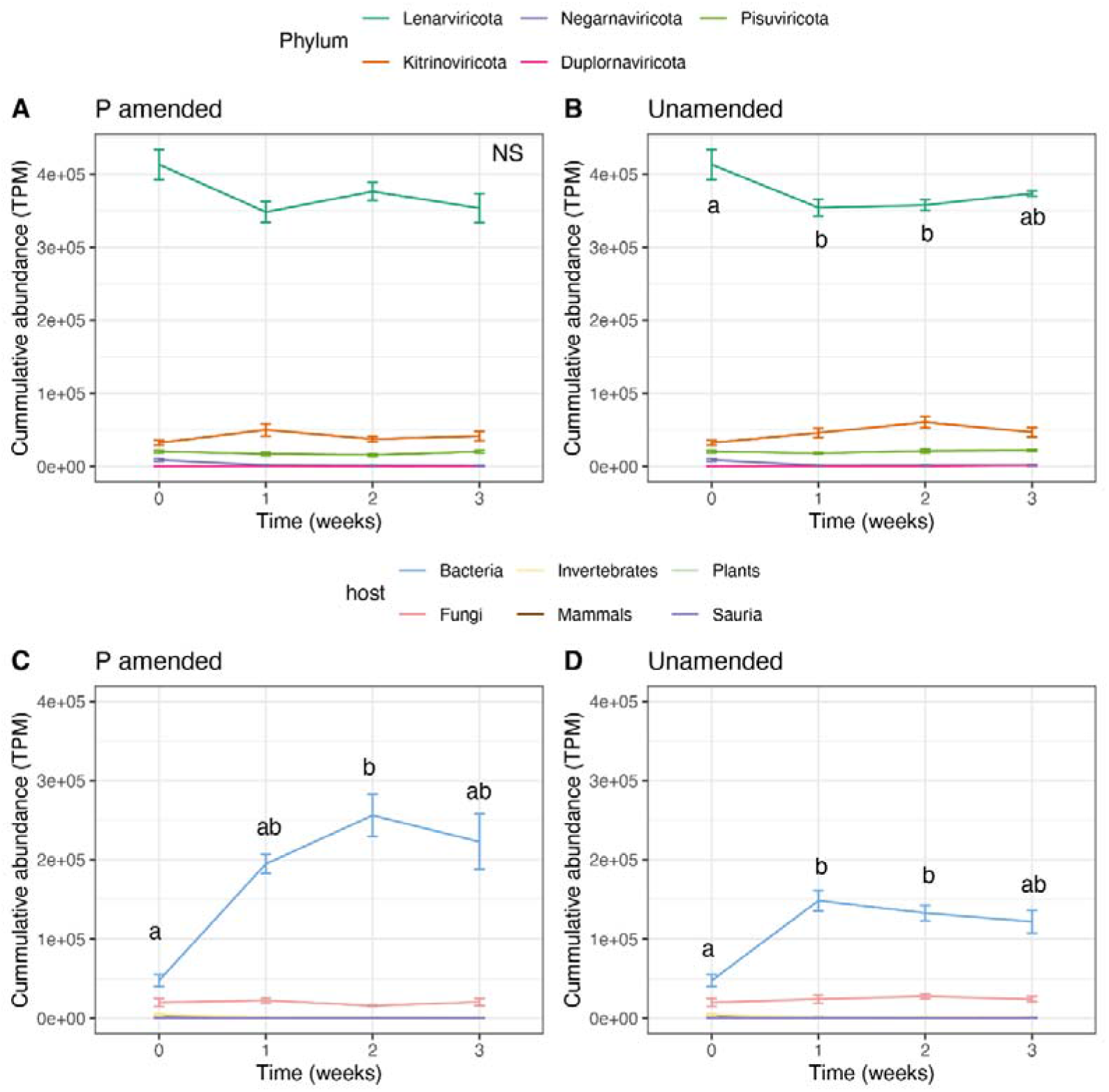
Temporal dynamics of RNA virus abundance following wet-up of a seasonally dry grassland soil. Cumulative abundance of soil RNA viruses over time in transcripts per million (TPM), with (A) or without (B) phosphate amendment. Colours represent viral phyla. Temporal dynamics of RNA viruses for which a host was predicted, plotted according to putative hosts, with (C) or without (D) phosphate amendment. Colours represent predicted host taxa. Error bars denote the standard error. Soil wet-up occurred immediately after T0 samples were collected. Different letters denote a significant change in abundance of *Lenarviricota* (A, B) or bacteriophages (C, D) at alpha = 0.1, n = 6 (Dunn test, Benjamini-Hochberg correction for multiple comparisons). NS = not significant.

The abundance of bacteriophages increased following wet-up, with a 5.4-fold increase in abundance when phosphate was added (*p* = 0.006; Fig. 3C), and a 3-fold increase in abundance when phosphate was not added (*p* = 0.01; Fig. 3D). Mycoviruses increased 3-fold during the first week and continued to increase when phosphate was not added; in contrast, there was no significant temporal pattern with phosphate amendment. Very few viruses had predicted eukaryotic hosts that were not fungi which decreased our statistical power to evaluate changes in their abundance.

To further investigate the bloom of phages post wet-up, we analyzed the amount of RNA representing the known phage groups *Leviviricetes* and *Durnavirales* (as well as other vOTUs predicted to infect bacteria over time), with or without phosphate amendment (Fig. 4A, 4B). In this experiment, the total amount of phage RNA increased dramatically within the first week (*p* < 0.1), and was more pronounced with phosphate amendment (*p* < 0.001; Fig. S11). This bloom can be interpreted as evidence of an active infection since an increase in the amount of viral RNA can only be achieved by viral replication; notably, the increase also corresponded with a significant increase in soil respiration (*p* = 0.03) (Fig. S12).

**Figure 4.**
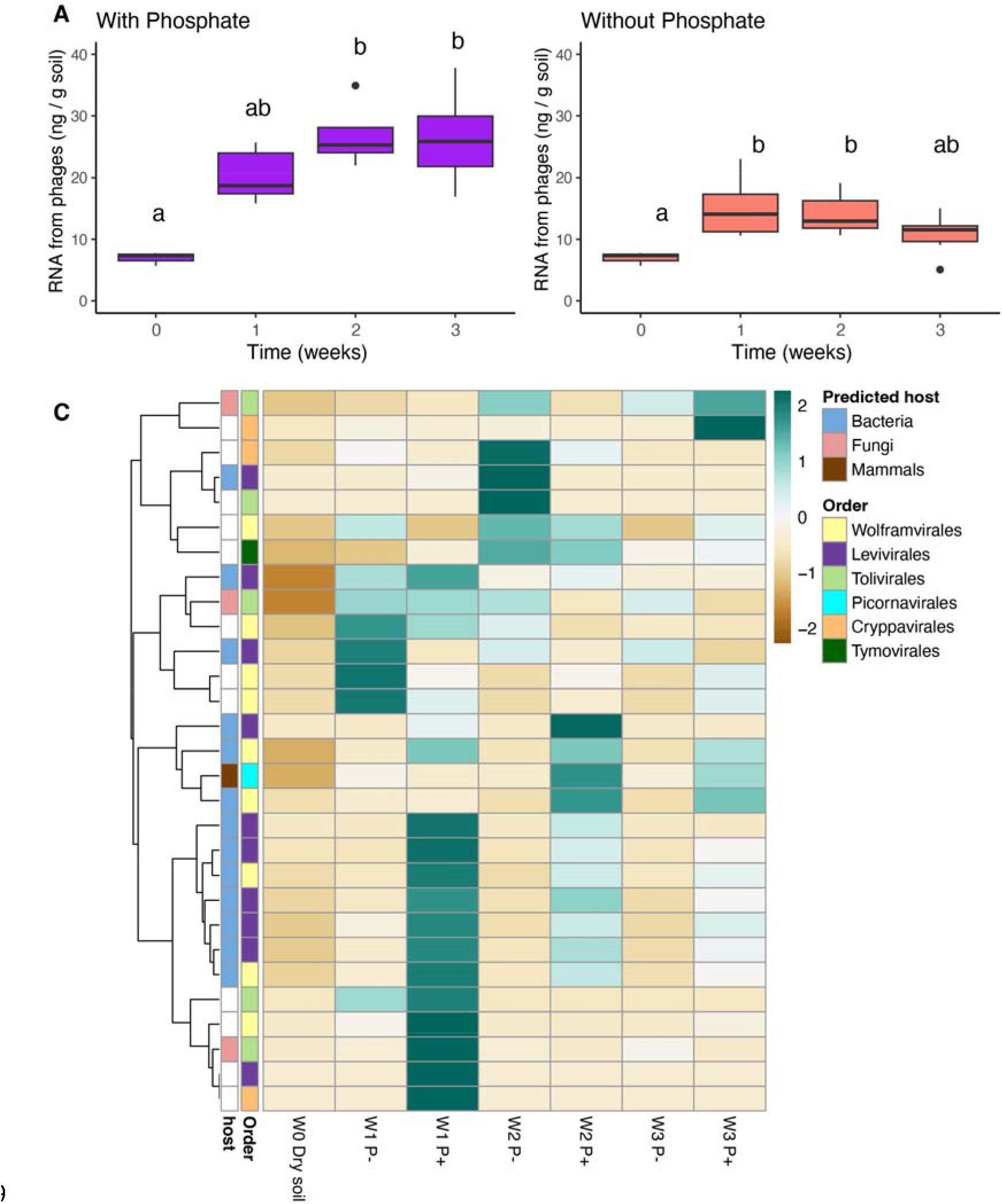
Bloom of bacteriophages and fungal viruses over time following soil rewetting. Boxplots representing the RNA abundance of bacteriophages with (A) and without (B) phosphate amendment. Based on the average genome size of cultured *Leviviricetes* (RefSeq, Nov 10, 2023), phage abundance was calculated by first determining the total RNA for phages, multiplying their normalized relative abundance by the proportion of virus-mapped reads and total extracted RNA per microcosm. Phage genome copies were then estimated by dividing this total RNA by the average genome weight. Finally, the number of bacterial hosts needed to generate this quantity of phage genomes were simulated, considering a range of genome copies per infected cell (virocell). (C) Heatmap showing vOTUs (rows) significantly up or downregulated by phosphate amendment (P+) (DESeq2, adjusted *p* < 10^−5^). The color scale denotes abundance Z-score normalized across rows, and the heat map is hierarchically clustered based on transcript abundance (TPM). Color bars to the left indicate the predicted host and taxonomy, with taxonomy assigned at the order level (Figs. S5–9).

Some of the vOTUs with significantly increased transcript abundance over the first week with phosphate amendment (adjusted *p* < 10^−5^) belong to *Wolframvirales* (8 of 29) rather than to the known phage class *Leviviricetes* (10 of 29), family *Cystoviridae* (0) or *Durnavirales* (0) (Fig. 4C). Many of the vOTUs that increased in response to phosphate amendment were predicted to infect bacteria (13 of 29), and three vOTUs of the order *Tolivirales* were predicted to infect fungi (Fig. 4C). With a less stringent *P* value requirement (adjusted *p* < 0.05) 80 vOTUs became more abundant one week after rewetting or with phosphate amendment, most of which were still *Wolframvirales* (24/80) and *Levivirales* (33/80) (Fig. S13).

The increase in phage abundance a week after soil rewetting (approximated by amount of viral RNA and normalised relative abundance of reads mapping to viruses (Tables S2, S4)) was used to estimate the number of cells containing active RNA phages (virocells) as a function of the average number of phage genome copies per cell and the average RNA phage genome weight (Table S4) (eq. 1). The genome weight is also affected by the number of strands in the phage genome, but as most of the phages identified belonged to the phylum *Lenarviricota*, which have single stranded RNA genomes, an average of one strand per phage genome was assumed. A range of 200–200,000 copies of phage genome per cell for the simulation (Eq. 1) was used, leading to a virocell count of 10^7^–10^10^ (for 200,000 and 200 copies per cell respectively; Fig S13).

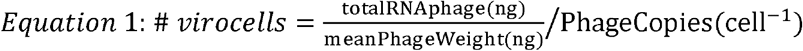

To explore whether the RNA viruses identified in our study might exhibit an extracellular phase, the open reading frame (ORF) annotations within their genomes was examined. Out of the 2,19 vOTUs, only a small subset (26 vOTUs) contained genes annotated as capsid or coat proteins, indicative of an extracellular phase and the potential to form extracellular virions (Tables S5, S6). These viruses, representing 0.8–3.5% of the total viral community (mean 1.3%), were primarily predicted to be bacteriophages, with two predicted to infect mammals. Notably, none of the phage genomes that were significantly enriched following phosphate amendment contained capsid or coat protein genes.

We characterized the bacterial and fungal communities in each microcosm to identify the diversity and temporal succession of potential hosts. Ordination of communities using phylogenetic distances (weighted unifrac) revealed that the overall bacterial community composition changed with phosphate amendment (Fig. 5A). In contrast, phages and mycoviruses were affected by phosphate amendment, time, and the interaction between the two factors (Fig. 5B).

**Figure 5.**
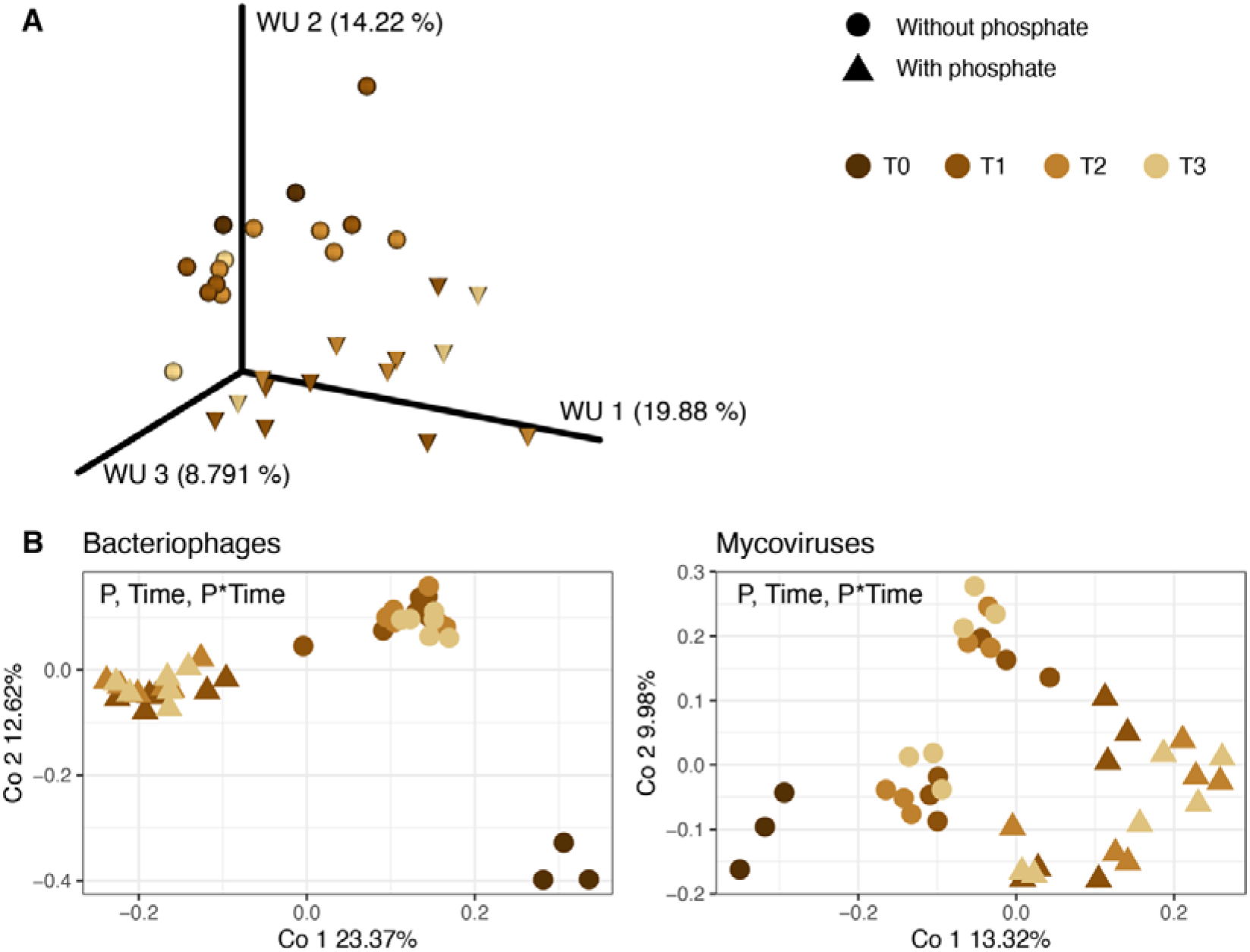
Beta diversity of grassland soil viral and putative host communities following wet-up and phosphate treatments. (A) Weighted unifrac (WU) distances of bacterial communities. (B) Ordination plots based on Bray-Curtis dissimilarity for RNA viruses (bacteriophage and mycoviruses). Time in weeks after wet-up (immediately after T0) is indicated by lighter brown colors. Treatments are denoted by shape: with phosphate (triangle) or without phosphate (circle) amendment. N = 6 per phosphate treatment and timepoint, with n = 3 at T0. Statistically significant effects (determined by PERMANOVA) are noted on each panel, in decreasing order of effect size. Phages: Phosphate *p* = 0.001, *R*^*2*^ = 22%, Time *p* = 0.001, *R*^*2*^ = 9%, P*Time *p* = 0.01, *R*^*2*^ = 5%. Mycoviruses: Phosphate *p* = 0.002, *R*^*2*^ = 5%, Time *p* = 0.001, *R*^*2*^ = 6%, P*Time *p* = 0.032, *R*^*2*^ = 6%.

After rarefying, we identified 3,914 bacterial and 629 fungal amplicon single variants (ASVs). Bacterial diversity based on the 16S rRNA gene significantly increased by the end of our experiment in unamended soils (Fig. S14). Fungal diversity was assessed with internal transcribed spacer region ITS2. As phylogeny based on the ITS2 region is not as robust as that of 16S rRNA, we present only phylogeny-independent analyses of fungal hosts. In both phosphate treatments, Shannon diversity decreased in the three weeks after wet-up (*p* = 0.08), with no significant differences between phosphate amended and unamended soils. These trends appear to be driven by changes in both richness and evenness (Fig. S14).

We analysed differential abundance and found 119 ASVs whose relative abundance was significantly lower in phosphate amended soil compared to unamended soil (*p <*10^−10^) (Table S7). Those ASVs belonged to *Actinobacteria, Acidobacteria, Planctomycetes, Alphaproteobacteria, Gammaproteobacteria, Chloroflexi, Deltaproteobacteria, Verrucomicrobia* and other phyla. All but two of these ASVs were undetectable before soil wet-up or with phosphate amendment. In unamended soil, only 26 of these ASVs (22%) were detected after one week. After two weeks 21/119 were detected (18%), and after three weeks 105/119 (88%).

## Discussion

In this study we used a replicated time-series of metatranscriptomes and amplicon analyses to investigate the response of RNA viruses and microbial communities to soil wet-up, and the added effects of phosphate amendment. Based on previous wet-up studies (Blazewicz et al., 2020), we hypothesized that soil rewetting would cause a ‘bloom’ in the microbial community (abundance, diversity and community composition of bacteria and fungi) and that this might in turn activate the RNA virus community. The community diversity indices of soil bacteria and fungi indicate that these host communities changed over the 3 weeks post-wetup and their structure was influenced by phosphate addition. Over this same period, soil respiration (CO_2_ flux) increased dramatically; particularly in the first week following rewetting, and at a more modest pace in the following two weeks. A dramatic shift in RNA virus community composition and the RNA abundance of bacteriophages occurred in the first week post wet-up, with relatively little change thereafter. The lack of change in the amount of RNA from phages in later time points may reflect either minimized replication of this class or zero-sum turnover. In a previous study of DNA viruses after a wet-up, the trajectory of the viral community over the first week appeared to be related to the particular viruses present in the field soil before wet-up (Nicolas et al., 2023). In contrast, our communities of RNA viruses clustered by amendment rather than by time, but the timeline of our current experiment was weekly rather than daily, and we also measured key changes to the RNA viral community during the first week. This suggests the major community assembly effects of the wet-up ‘bloom’ occur within the first week after wet-up, while the effects of phosphorus limitation on hosts and RNA viruses can persist for longer.

Due to their high stoichiometric demand for phosphorus, we have good reason to expect that rapidly growing viral populations might cause localized phosphorus limitation (Kuzyakov and Mason-Jones, 2018; Tong et al., 2023). Unlike their hosts, viral particles (virions) have a minimal amount of lipids and membranes and are composed mainly of proteins and nucleic acids which are comparatively rich in phosphorus; therefore, they have a lower C:N:P ratio than cells (Jover et al., 2014). If microbial necromass produced during a viral proliferation event has a reduced amount of phosphorus, this could lead to phosphorus limitation for organisms feeding on the necromass (Kuzyakov and Mason-Jones, 2018).

However, contrasting evidence suggests viral proliferation could alleviate phosphorus limitation via lysis of host cells, releasing organic material into the immediate environment, or that bacteria may even predate on phages during phosphorus limiting conditions (Godon et al., 2021; Tong et al., 2023). In our study, we hypothesized that added phosphate would stimulate RNA virus infection by alleviating phosphorus limitation.

We observed a clear effect of phosphate amendment on the community of RNA viruses. Phosphate amendment significantly altered RNA viral community composition within the first week after rewetting, and phosphate amended versus unamended communities remained distinct for up to three weeks. While the effect of soil wet-up appears to wane one week after wet-up, the differences in viral community structure between amended and unamended soil were maintained throughout three weeks. While this provides evidence of phosphorus limitation, it is unclear whether it stems from viral proliferation. The cumulative metabolic rate of potential hosts (proxied by respiration) as well as bacterial and fungal community composition were unaffected by phosphate amendment, indicating that phosphorus is not the limiting factor for the host community, only to viruses. However, when accounting for host phylogeny, bacterial communities were affected by phosphate amendment, implying that while the community as a whole is not phosphorus-limited, specific taxonomic clades are. The estimated amount of viral RNA extracted from phosphate amended soil was higher compared to unamended soil for three weeks following rewetting, supporting our hypothesis and implying that the amendment triggered viral proliferation.

The vast majority of RNA viruses we identified belong to the phylum *Lenarviricota* (ssRNA+), which are predicted to infect bacteria or fungi. This is consistent with community composition of RNA viruses from a nearby site during plant growth, as well as from grasslands in England and Kansas, USA—although the relative abundance of RNA phages we found was somewhat lower (ca. 20%) than in others studies (50% on average) (Starr et al., 2019; Wu et al., 2021; Hillary et al., 2022). This difference could be due to the type of soil or environmental conditions (e.g., presence or absence of plants, soil moisture) which supports different host communities. Since our study was performed on dry soil without plants, we expected the majority of viruses to infect microorganisms such as fungi and some bacteria that can survive dry summer conditions. Fungal viruses (mycoviruses) are particularly likely to survive during the dry season as they do not have a free-living, extracellular phase (Hough et al., 2023).

RNA bacteriophages significantly proliferated after soil wet-up, more so in phosphate amended soil. Most of the phage vOTUs that became more abundant with phosphate amendment were classified as *Leviviricetes* (the largest clade of RNA phages) and *Wolframvirales*. RNA phages have only eleven representatives in pure culture with curated complete genomes (RefSeq Dec 6^th^ 2023). All eleven are ssRNA phages that belong to the class *Leviviricetes* (formerly phylum *Leviviricota*) and have genomes ranging in size between 3,400–4,300 bases. Moreover, these phages were isolated on bacteria that are human pathogens or part of the human microbiome, belonging to the phyla *Proteobacteria* or *Pseudomonadata* (Zinder, 1980; Klovins et al., 2002). Recently, a study based on bioinformatic characterization of RNA viruses proposed that the dsRNA order *Durnavirales* is comprised of phages (Neri et al., 2022). However, this order also has no cultured representatives. Therefore, we suggest that future work should concentrate on isolation of RNA phages from different environments within environmentally relevant hosts to characterize the effects of RNA phages on biogeochemical cycling.

The importance of microbial mortality in soil has been highlighted in recent reports showing that a notable amount of persistent, mineral-associated soil organic matter is comprised of microbial necromass (Liang et al., 2019; Sokol et al., 2019, 2022; Angst et al., 2021). Viruses are a potentially important driver of microbial mortality in soil, as estimated in marine environments (Boras et al., 2009; Vaqué et al., 2017) and described as the ‘viral shunt’ (Wilhelm and Suttle, 1999; Suttle, 2007). Up to 40% of soil bacteria are thought to contain lysogenic phage that can become lytic if induced by a dramatic change in resource availability or environmental conditions (Fuhrman and Schwalbach, 2003; Williamson et al., 2007), such as appearance of a newly growing root (Starr et al., 2019, 2021), or a rewetting event (Nicolas et al., 2023; Santos-Medellín et al., 2023). Nicolas et al.’s (2023) recent study on the succession of DNA viruses following soil wet-up identified a temporal increase in DNA extracted from the viral size-fraction. This increase was interpreted as lytic infection of host cells, particularly bacteria, leading to an increase in virions and up to 46% of microbial cell deaths attributed to viral activity in the week after soil wet-up (Nicolas et al., 2023).

We estimated replication of RNA phages via the increasing amount of phage RNA and demonstrated their proliferation over the first two weeks following soil wet-up. However, because our data was drawn from bulk soil metatranscriptomes rather than RNA from a viral targeted metatranscriptome (the viral size fraction), the existence of RNA virions in the environment cannot be inferred. While cultured RNA phages are known to be lytic, many other RNA viruses do not have an extracellular phase in their life cycle, and do not kill their host but rather infect it chronically (Nuss, 2011). Moreover, no more than 3% of the phage genomes assembled contained a coat or capsid protein. Culturing viruses by plaque assays creates an inherent bias towards lytic viruses and precludes us from culturing non-lytic RNA viruses. Therefore, we used burst sizes of cultured RNA phages to estimate the number of infected cells (virocells) rather than microbial mortality. Whether or not free virions of the RNA phages identified in this study exist, these phages would certainly affect their hosts dramatically, as active infections can rewire host metabolism and redirect resources from the cell towards viral replication. The number of bacteria per gram of soil has been estimated to range between 10^7^–10^10^ (Forterre, 2013; Williamson et al., 2017). The lowest number of virocells we extrapolated, based on an average of 200,000 copies of phages genomes per virocell, was on the order or 10^7^ cells per gram of soil, potentially accounting for up to 100% of bacteria. Therefore, we suggest the RNA phages may have an important role in shaping the bacterial community following soil wet-up, with at least a comparable impact to that of DNA phages.

Our study aimed to describe the diversity and abundance patterns of RNA virus communities in Mediterranean grassland soil during the three weeks following rewetting, while testing whether viral and host communities were limited by phosphorus availability. We found that some RNA viruses were phosphorus-limited, while host communities were not. Notably, RNA phages bloomed in response to phosphorus amendment, potentially infecting a significant proportion of bacteria in rewetted soil. The most significant shifts in RNA viral communities occurred within the first week post-wet-up, suggesting a normalization period following the disturbance. Unlike prior studies on DNA viruses, which indicated that community trajectories were shaped by pre-existing viral populations, our results showed that RNA viral communities clustered more by phosphorus amendment than by time. Phosphate amendment significantly altered the RNA viral community, with differences persisting for three weeks, indicating that phosphorus availability impacts viral, not host, communities and shapes their composition. The majority of RNA viruses identified belonged to the phylum *Lenarviricota*, with RNA bacteriophages significantly proliferating after rewetting. Although we could not directly measure viral-induced mortality, we estimated that a substantial proportion of bacterial cells were likely infected by RNA phages, suggesting these viruses play a critical role in shaping bacterial communities. Overall, our findings provide important information on RNA virus communities and their host interactions in soil ecosystems, particularly their responses to environmental changes and nutrient amendments, indicating that RNA phages may influence biogeochemical cycling in a manner similar to DNA phages.

### Code availability

R code used in this study is available on https://github.com/ellasiera/RNA_virus_wetup_2023.

## Acknowledgments

We acknowledge that Hopland Research and Extension Center sits on the traditional, unceded land of the Pomo Indians. Research conducted at Lawrence Livermore National Laboratory took place on the territory of xučyun (Huichin), the ancestral and unceded land of the Chochenyo-speaking Ohlone people. We thank Christina Ramon and Julie Johnston for help in harvesting the microcosms and Jessica Wollard for help with extracting and sequencing DNA for the amplicon data. We thank Andy Millard for providing a courtesy review of the manuscript.

## Funding

The work was supported by a Lawrence Livermore National Laboratory, Laboratory Directed Research & Development grant (21-LW-060) to GT and by LLNL’s U.S. Department of Energy, Office of Biological and Environmental Research, Genomic Science Program “Microbes Persist” Scientific Focus Area (#SCW1632). ETS was supported by a Marie Skolodowska-Curie postdoctoral fellowship “Divobis”. Work conducted at LLNL was conducted under the auspices of the US Department of Energy under Contract DE-AC52-07NA27344.

## Author contributions

GT conceptualized the study, acquired funding, ran the experimental procedures, supervised, and revised the paper. ETS performed data curation, formal data analysis, visualization, writing and revising. GME assisted with experimental procedures and writing and editing. JAK, GWN, CH, EEN, SJB and JPR assisted with experimental procedures, supervised data analyses, edited and revised the manuscript.

## Notes

### Competing Interest Statement

The authors have declared no competing interest.

