## Supplemental Figures 1-14 for "Phosphate amendment drives bloom of RNA viruses after soil wet-up"

**Figure S1:** Number of microcosms a vOTU appears in (N=3300). A third of vOTUs appear only in one microcosm.

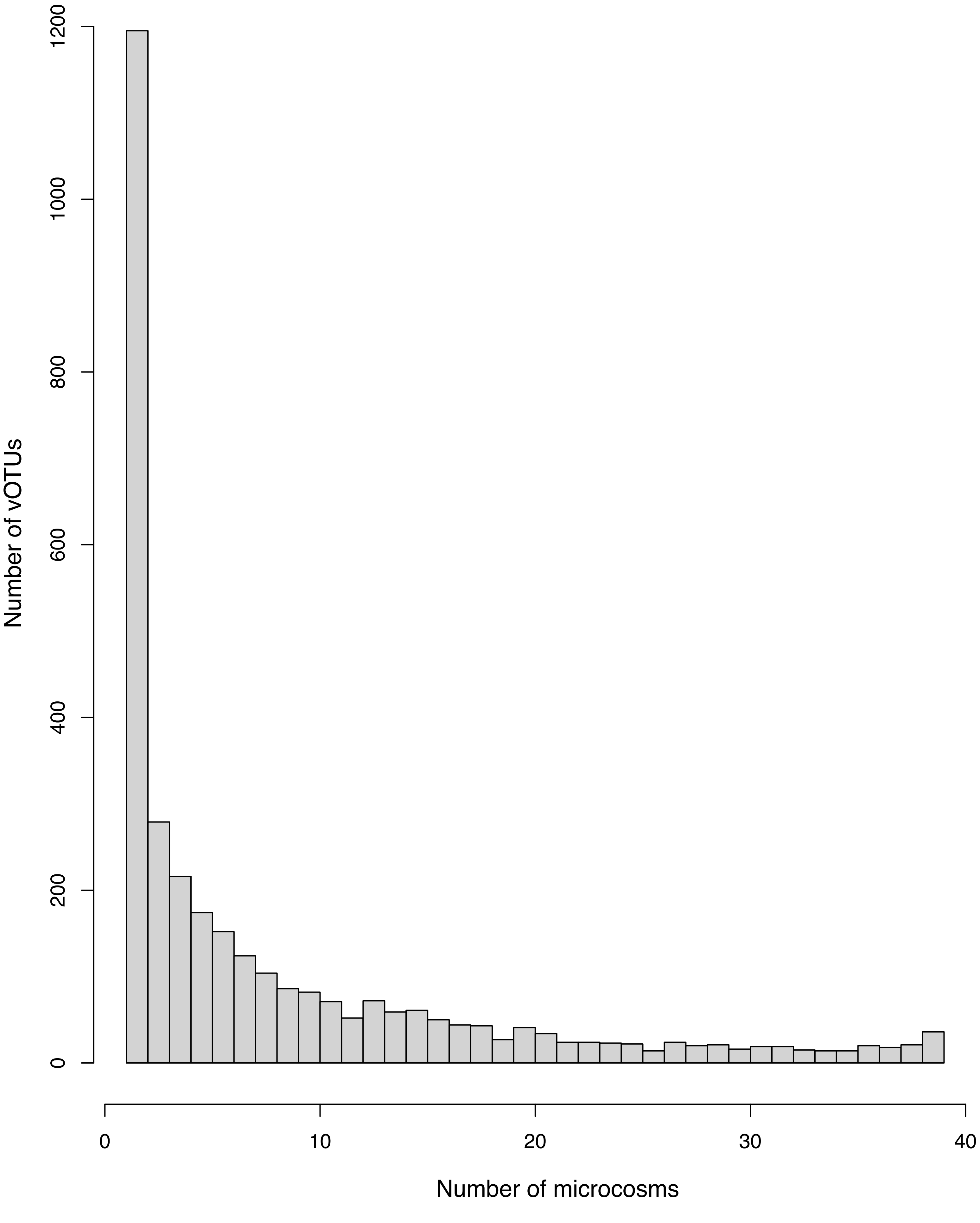

**Figure S2**: Distribution of rank of vOTUs belonging to the most abundant viral classes across any number of microcosms. Overall, the taxonomic distribution of vOTUs appearing in X samples is independent of X (1<X<39).

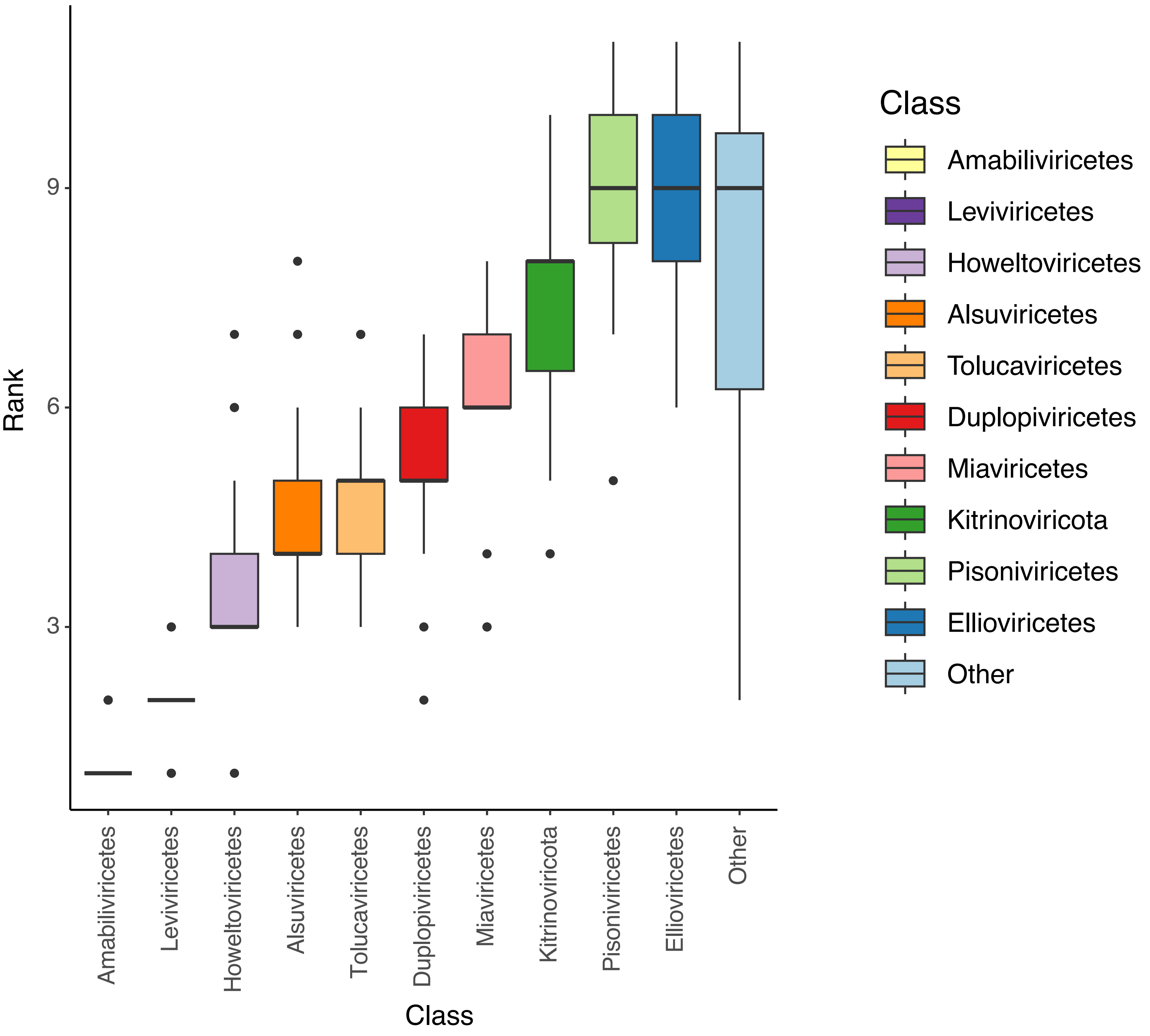

**Figure S3**: Number of unique and shared vOTUs between replicates over time followings soil wetup, with/ without phosphate amendment.

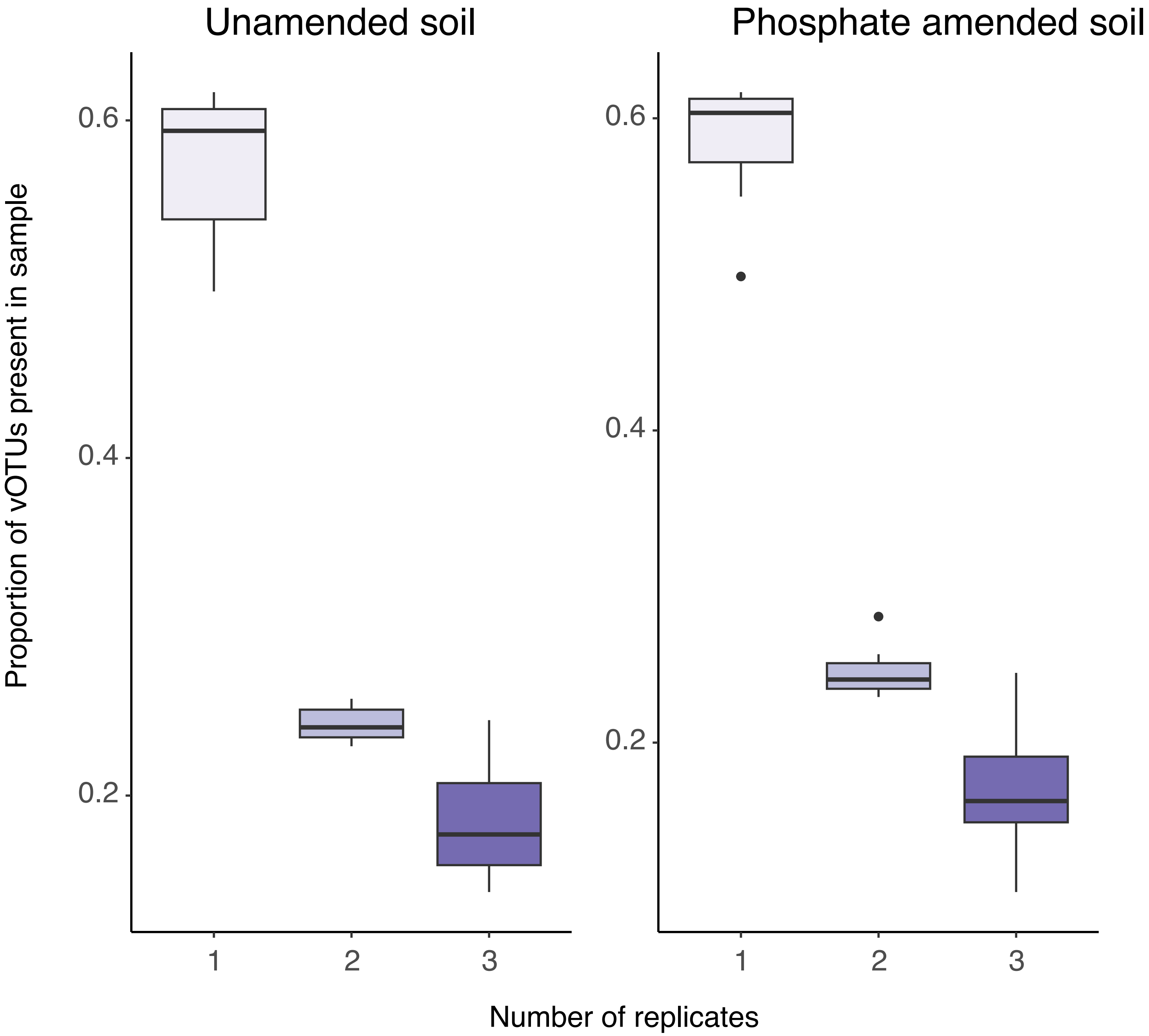

**Figure S4:** Taxonomy and transcripts per million (TPM) of RNA viral operational taxonomic units (vOTUs) that were present only in dry soil, or only in phosphate-amended soil, or only in unamended soil following wetup. As these are RNA viruses, transcription level may represent abundance and/or activity.

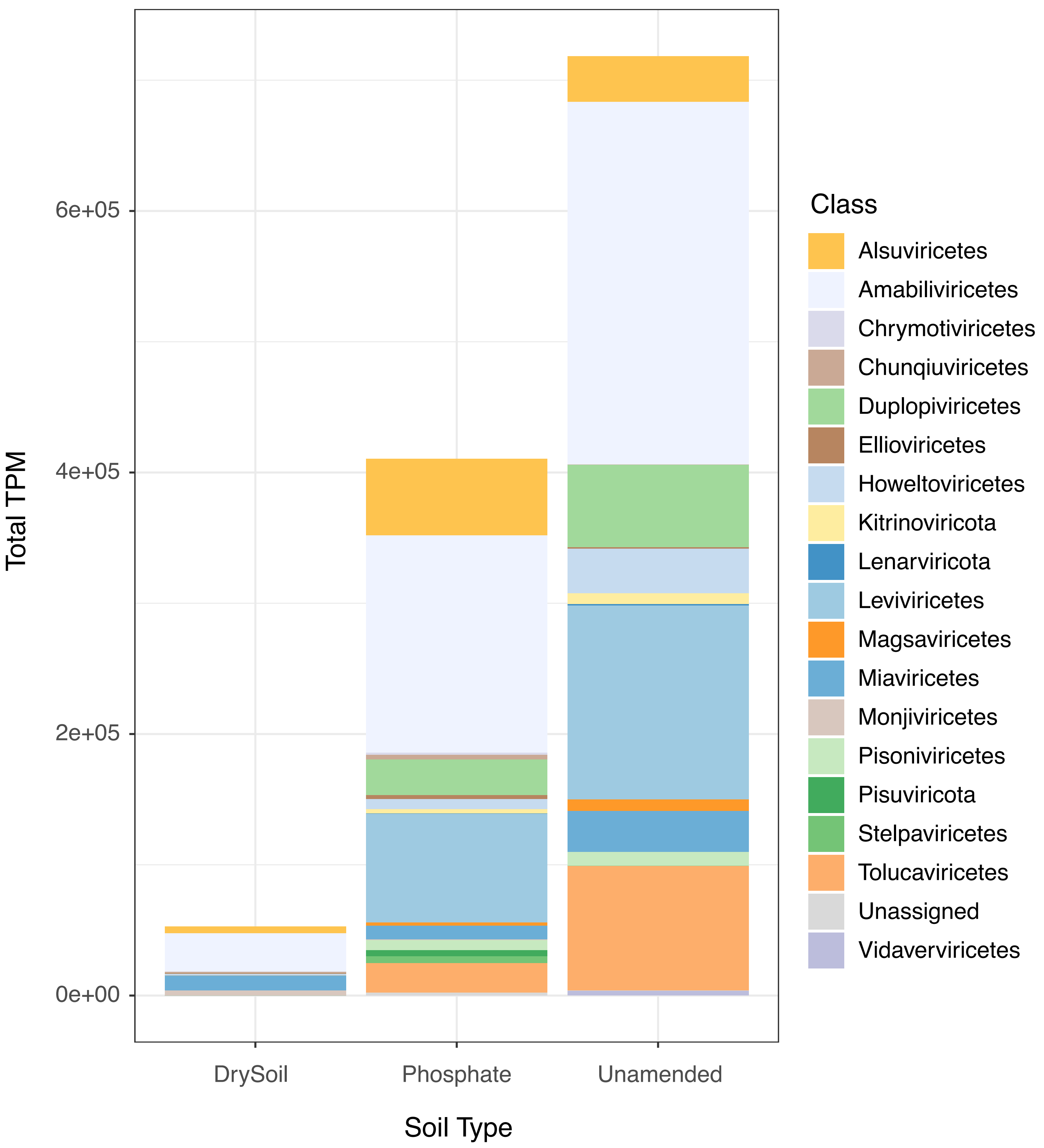

**Figure S5**: Phylogenetic tree of RdRp protein sequences from phylum Negarnaviricota. Black leaf labels indicate reference sequences and red leaf labels mark sequences from this study. The inner concentric circle displays taxonomy at the order level, and the outer circle shows known hosts. The tree is based on a trimmed MAFFT alignment and was built in IQtree2.

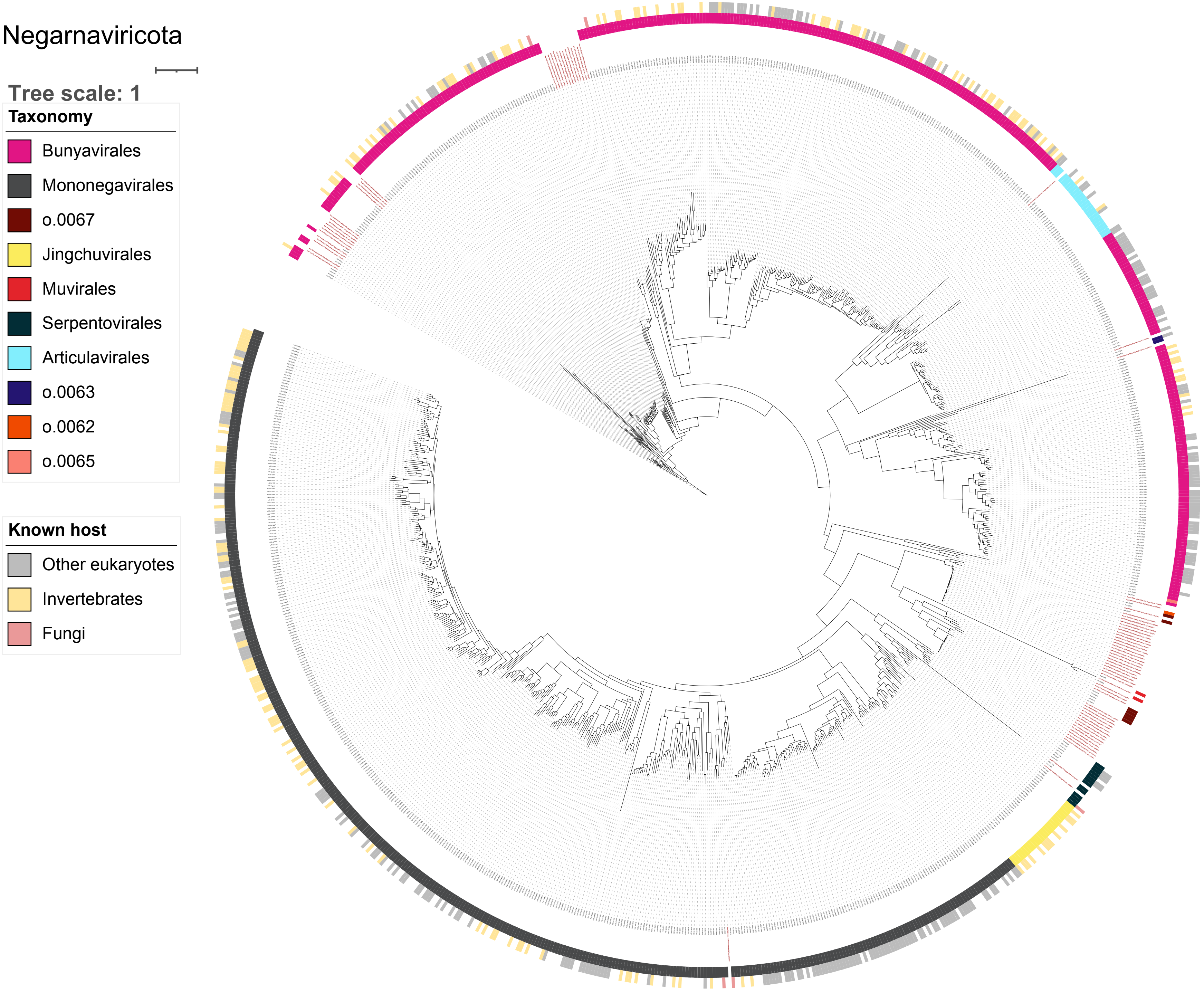

**Figure S6**: Phylogenetic tree of RdRp protein sequences from phylum Kitrinoviricota. Black leaf labels indicate reference sequences and red leaf labels mark sequences from this study. The inner concentric circle displays taxonomy at the order level, and the outer circle shows known hosts. The tree is based on a trimmed MAFFT alignment and was built in IQtree2.

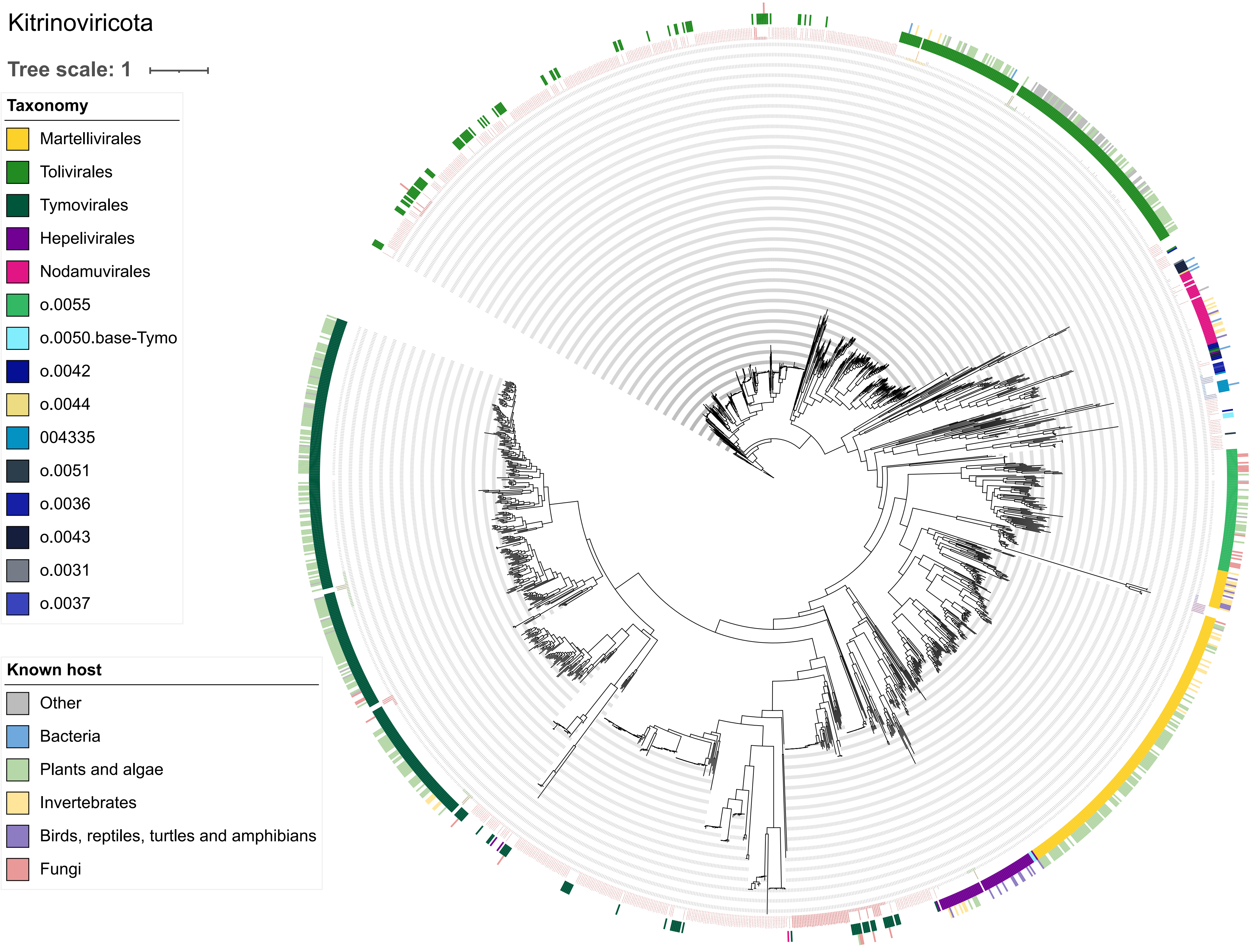

**Figure S7**: Phylogenetic tree of RdRp protein sequences from phylum Pisuviricota. Black leaf labels indicate reference sequences and red leaf labels mark sequences from this study. The inner concentric circle displays taxonomy at the order level, and the outer circle shows known hosts. The tree is based on a trimmed MAFFT alignment and was built in IQtree2.

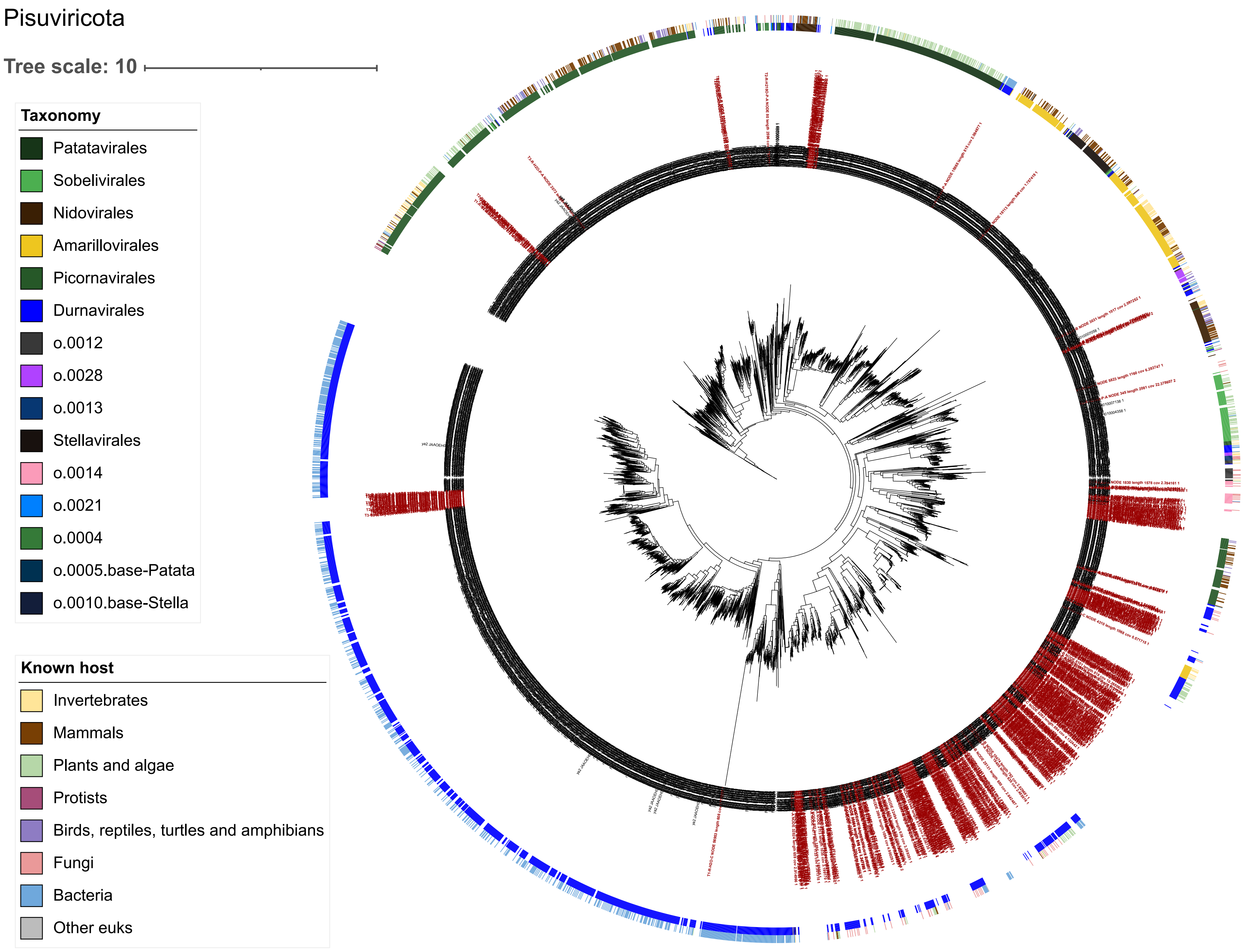

**Figure S8**: Phylogenetic tree of RdRp protein sequences from phylum Duplornaviricota. Black leaf labels indicate reference sequences and red leaf labels mark sequences from this study. The inner concentric circle displays taxonomy at the order level, and the outer circle shows known hosts. The tree is based on a trimmed MAFFT alignment and was built in IQtree2.

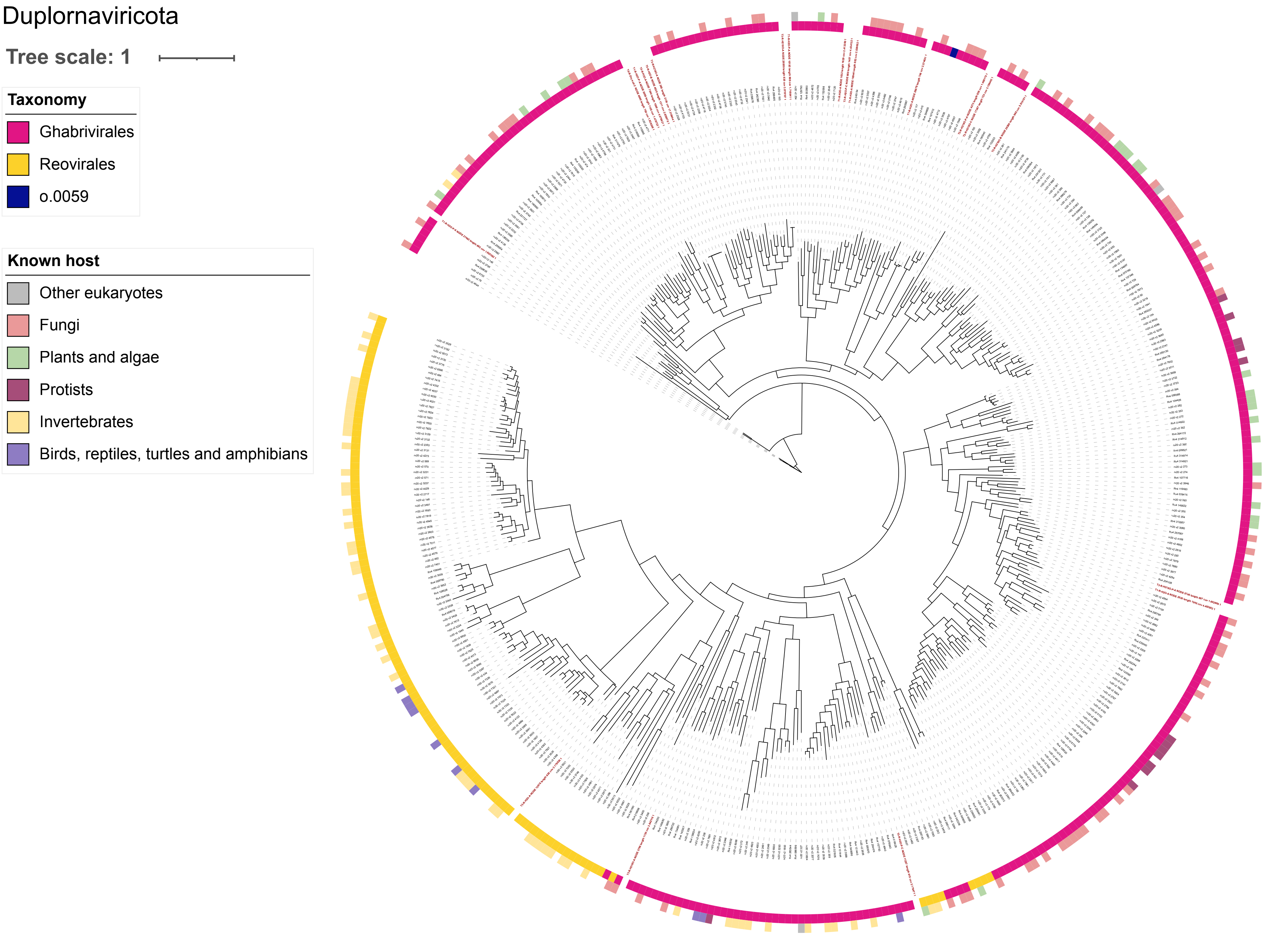

**Figure S9**: Phylogenetic tree of RdRp protein sequences from phylum Lenarviricota. Black leaf labels indicate reference sequences and red leaf labels mark sequences from this study. The inner concentric circle displays taxonomy at the order level, and the outer circle shows known hosts. The tree is based on a trimmed MAFFT alignment and was built in IQtree2.

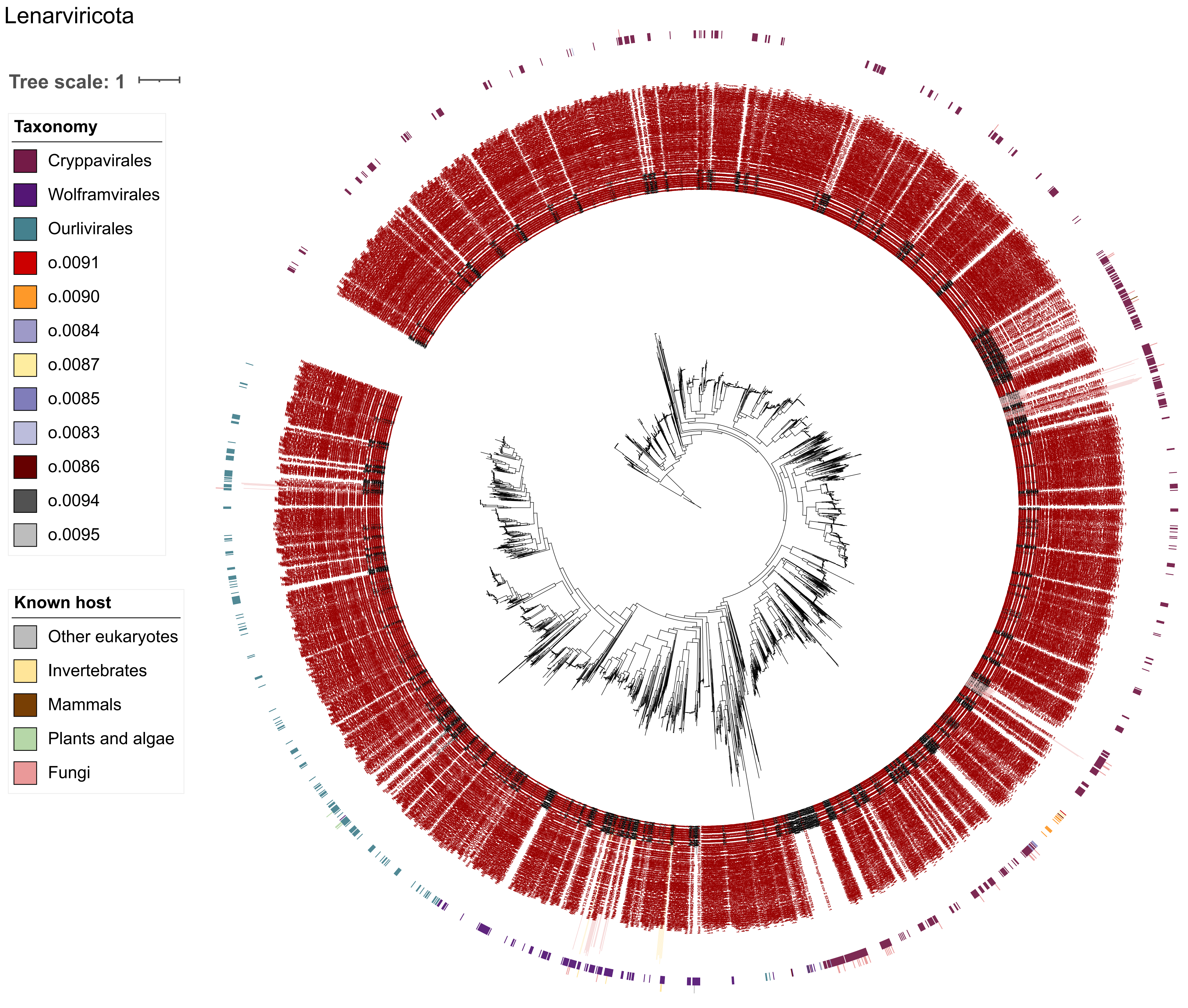

**Figure S10:** Temporal abundance of RNA virus phyla after soil rewetting with or without phosphate amend-ment.

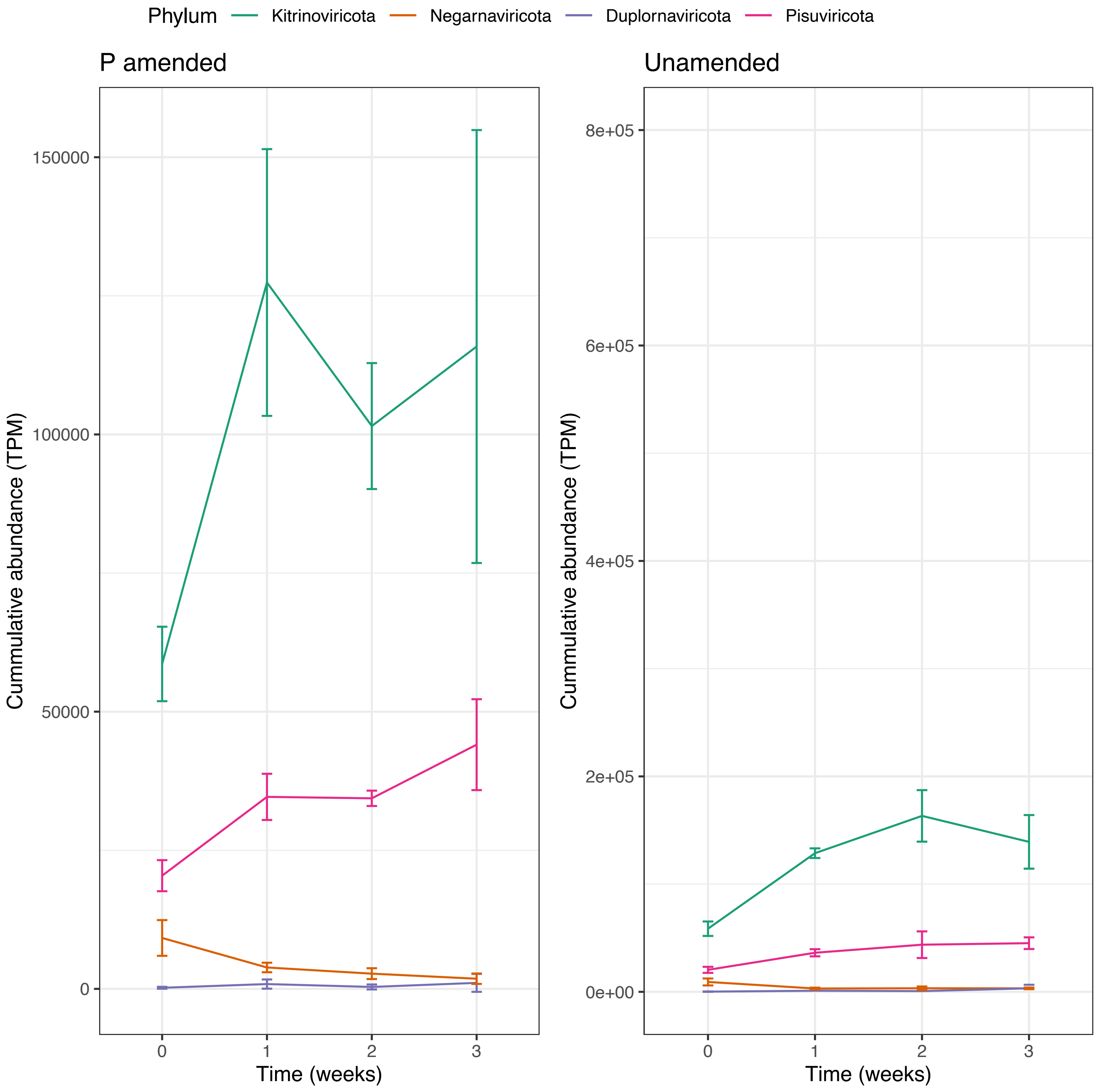

**Figure S11**: RNA Phage bloom following soil rewetting with or without phosphate amendment. Amount of RNA from phages based on their relative abundance out of the entire RNA virus community multiplied by the relative abundance of reads mapped to viral contigs out of total RNA with (A) and without (B) phosphate amendment during wet-up. The number of bacteria infected by RNA phages was simulated by the change in phage RNA between consecutive time points divided by the average weight of an RNA phage genome based on curated public data and divided again by the simulated number of phage copies per cell with(C) and without (D) phos-phate amendment.

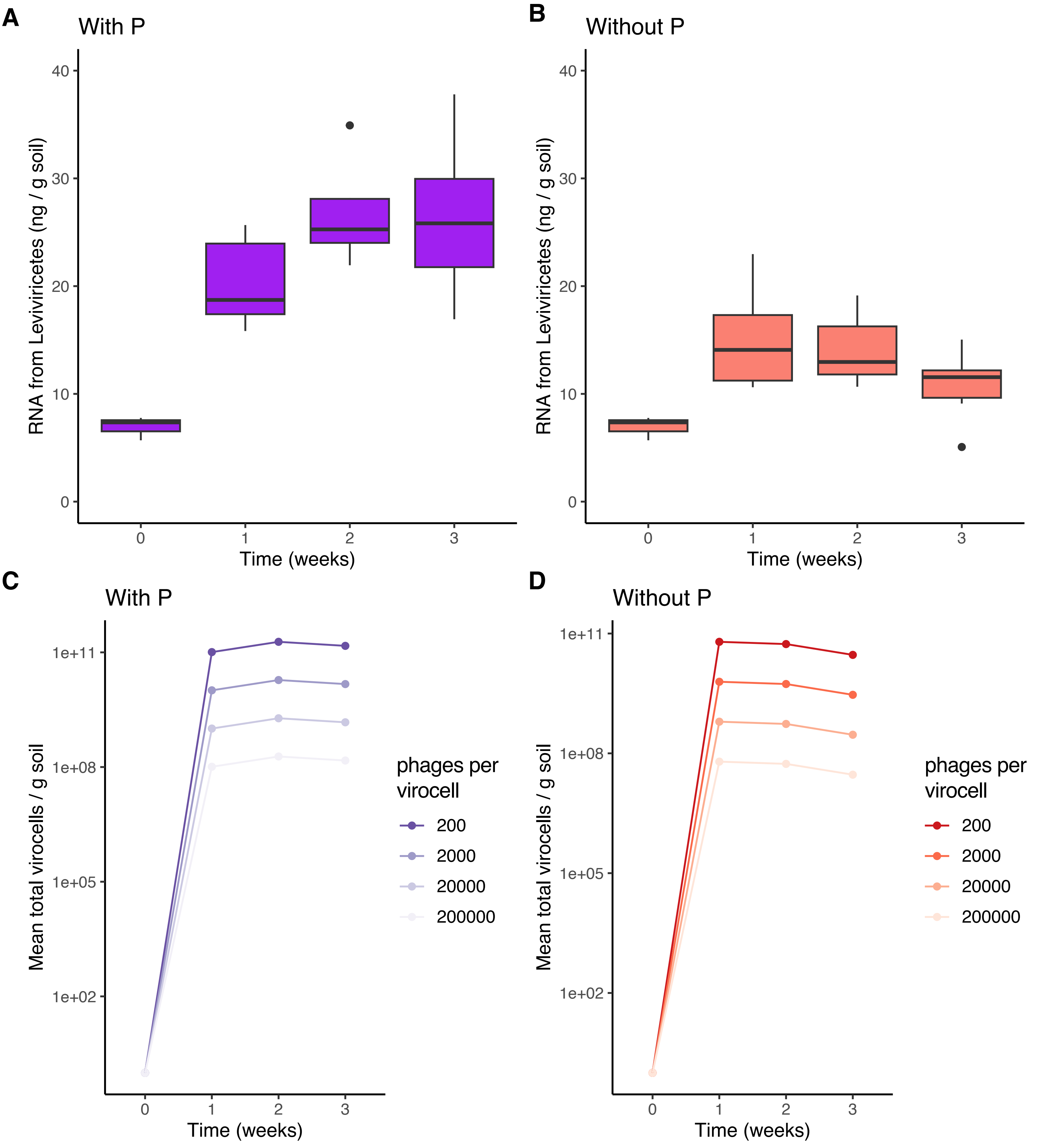

**Figure S12**: Respiration before (0) and after soil rewetting measured from microcosm headspace (N=6).

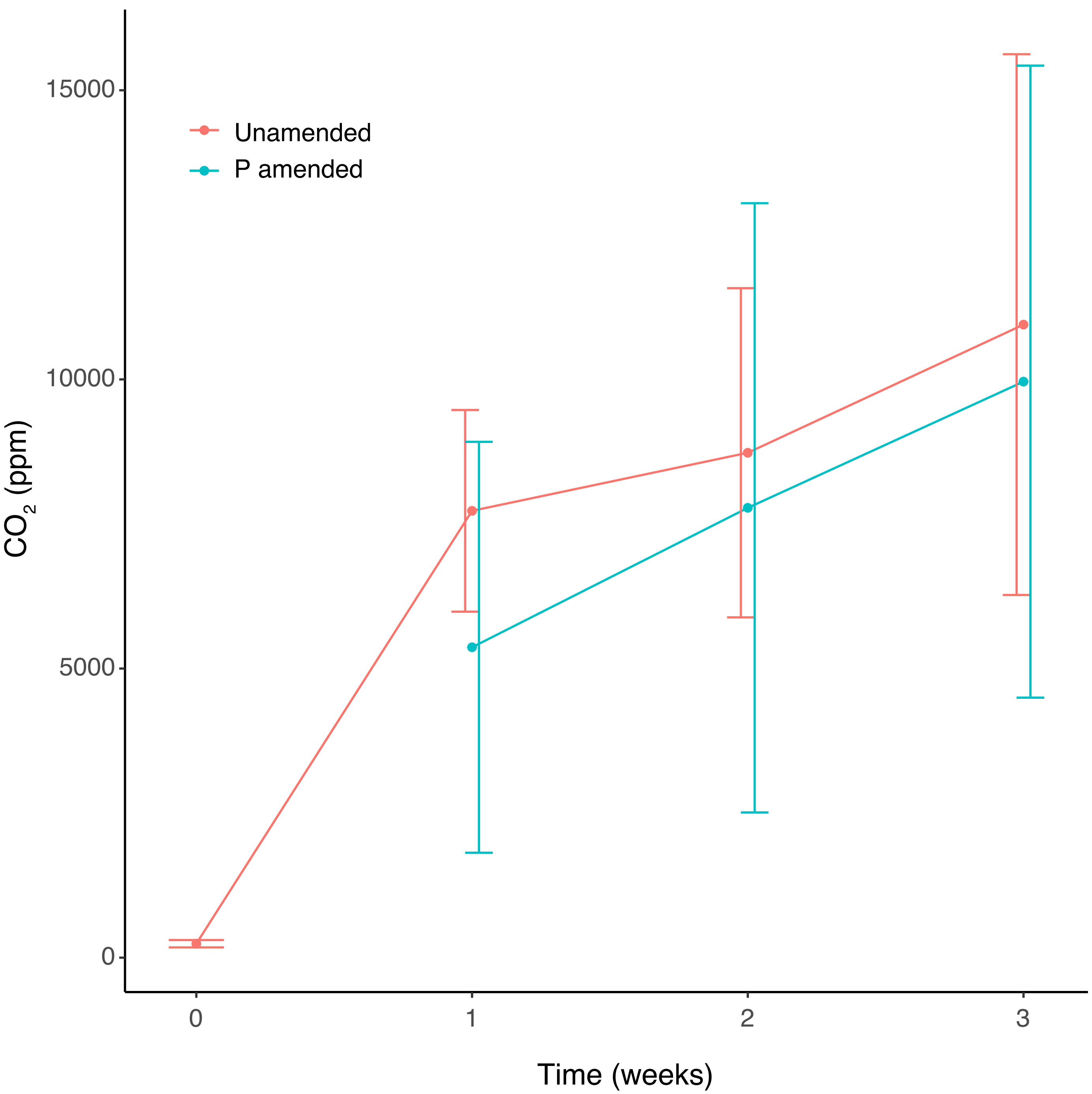

Figure S13: Transcript abundance (TPM) of vOTUS whose abundance increased significantly (p <=0.05, n=80) over the first week after soil rewetting or with phosphate amendment. Abundance is Z-score normalized per row. Hierarchical clustering, host prediction and taxonomy are presented to the left of the TPM heat map. P+ denoted phosphate amendment, and P- unamended soil.

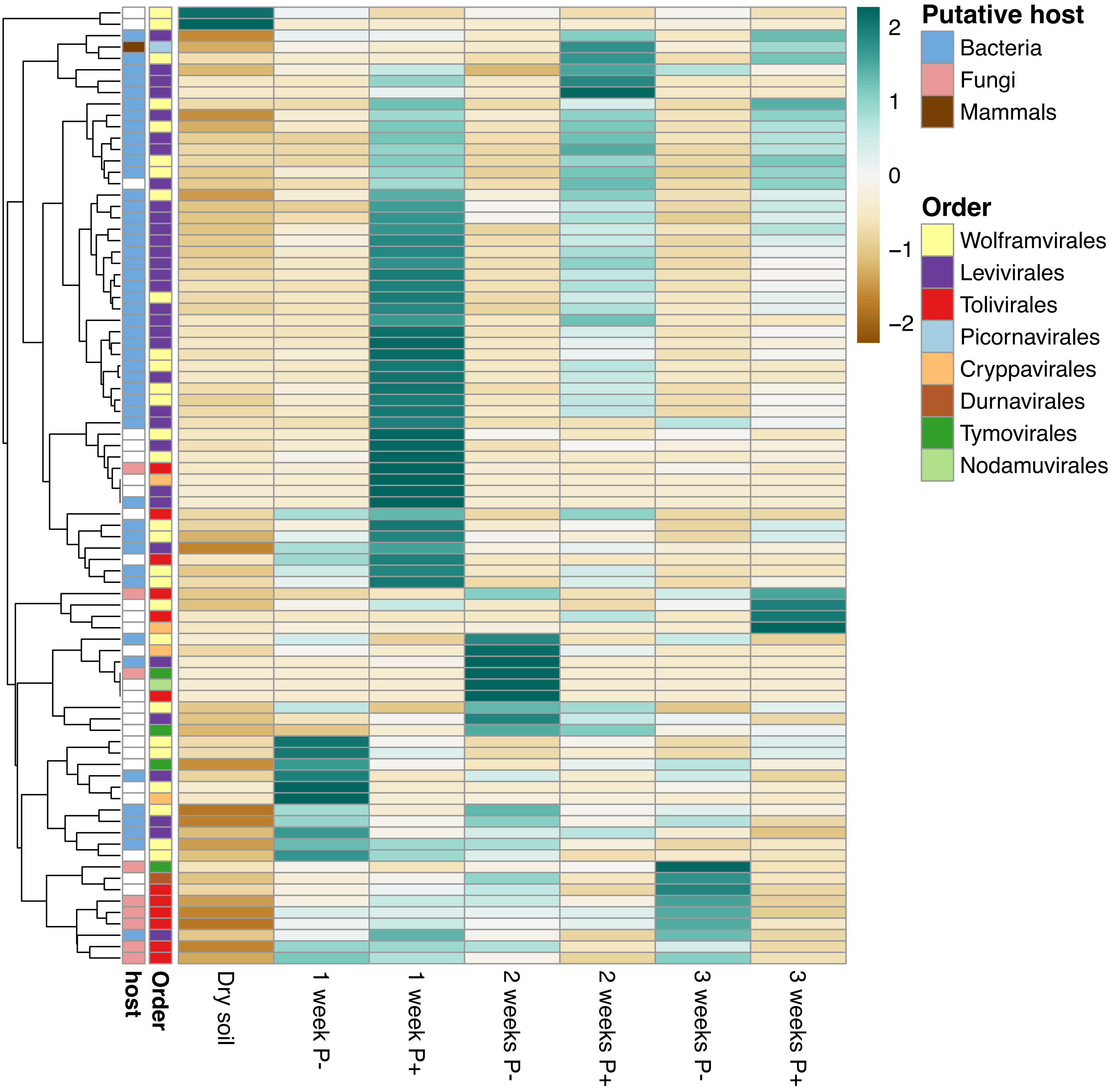

**Figure S14:** Diversity, richness (species number) and evenness of (A) fungal and (B) prokaryotic communities before and after soil wetup with or without phosphate amendment.

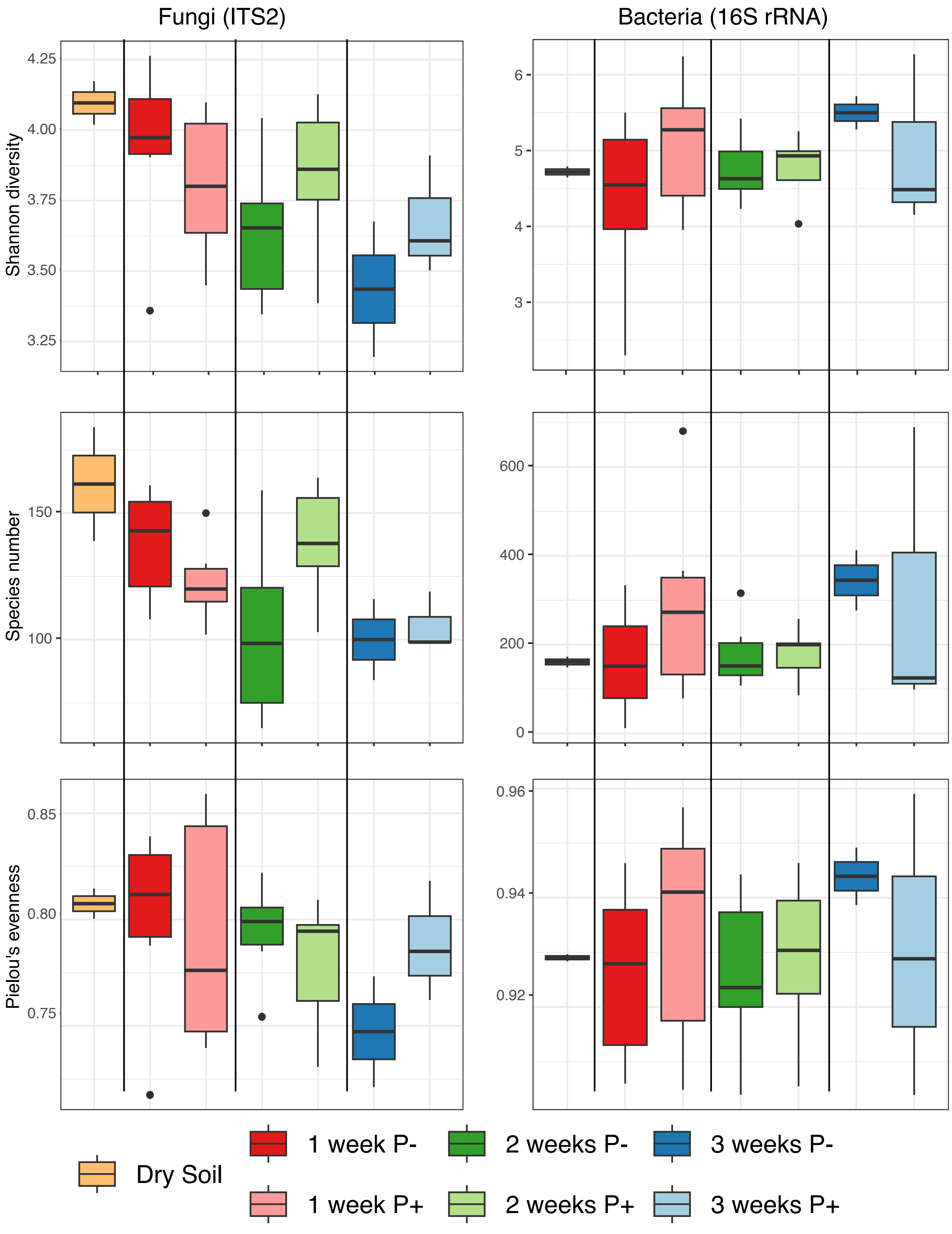
